# Single cell mapping reveals phenotypic, immunometabolic and functional heterogeneity among age-associated B cells in systemic lupus erythematosus and aging

**DOI:** 10.64898/2026.11.23.767014

**Authors:** Emmanouil Stylianakis, Cedric Bissinger, Julia Weinmann-Menke, Nadine Hövelmeyer

## Abstract

**Introduction:** Age-associated B cells (ABCs) comprise an antigen-experienced memory B-cell population that expands during aging but has also been implicated in autoimmune diseases, particularly systemic lupus erythematosus (SLE). Cell immunometabolism has emerged as a key determinant of immune-cell function, linking metabolic adaptation to both protective and maladaptive immunity or pathological inflammation, thereby providing a platform for therapeutic intervention. This study aims to comprehensively profile the functional alterations that accommodate ABC expansion, emphasizing metabolic reprogramming within the B-cell compartment in SLE, particularly with regard to ABCs, in order to identify niche metabolic pathways utilized by ABCs that may represent useful therapeutic targets.

**Methods:** An exploratory analysis was conducted using publicly available single-cell RNA sequencing datasets to phenotypically and functionally characterize ABC populations in patients with SLE and lupus-prone MRL.lpr mice, using a broad array of community tools. We further conducted SCENITH-based metabolic profiling to assess metabolic differences among splenic follicular (FO), marginal zone (MZ) and CD21^-^CD23^-^ double negative (DN) B cells, in the murine SLE-like MRL.lpr model, in which we tested the ability of B cell subsets to generate protein output, following exposure to specific metabolic inhibitors (2-deoxyglucose and oligomycin). In addition, we cross-examined CD21⁻CD23⁻ DN B cells in the native context of aging by employing flow cytometry-based analyses of their redox status (NADH/FAD autofluorescence) and mitochondrial function (MitoTracker and TMRE staining) to investigate age-associated metabolic programming.

**Results:** Single-cell transcriptomic analyses revealed substantial immunophenotypic and metabolic heterogeneity within the ABC compartment in SLE. Subclustering of ABCs revealed three separate and distinct ABC subsets, encompassing CD68⁺ ABCs, NKG7⁺ ABCs, and AREG⁺ ABCs, differing in their respecrive transcriptional, functional, and immunometabolic signatures. The CD68⁺ ABC subset demonstrated signs of enhanced cytotoxicity, phagocytosis, lysosomal activity, and increased antigen-processing capabilities. The CD68⁺ and NKG7⁺ ABC populations exhibited increased inflammatory pathway activity, including pattern-recognition receptor signalling and NLRP3 inflammasome-associated activity, while the AREG⁺ ABC subset displayed signs of senescence, moderation, exhaustion, or anergy. hdWGCNA identified transcriptional modules associated with acetyltransferase activity, suggesting a potential link between metabolic reprogramming and epigenetic remodelling through epimetabolite availability, in particular acetyl-CoA. ABCs demonstrated, in general, coordinated amplification of glycolysis, oxidative phosphorylation, and lipid utilization, with substantial intragroup metabolic heterogeneity among the three ABC subsets. The CD68⁺ ABCs emerged as the most hypermetabolic subset. In parallel, SCENITH-based metabolic comparison of CD21⁻CD23⁻ DN B cells between mildly diseased male and severely diseased female MRL.lpr mice revealed increased utilization of alternative non-glucose substrates and enhanced glycolytic activity, but reduced overall protein synthesis capacity, indicating altered allocation of metabolic resources. Further interpretation of the transcriptomic analysis suggests that proteasomal activity could explain the lower protein levels in female MRL.lpr mice, as measured by SCENITH. In addition, in silico drug prediction pointed towards statins and proteasome inhibitors as potentially useful pharmacological agents for blocking cholesterol metabolism and protein degradation, respectively. Finally, the analysis of aged mice also revealed metabolic and redox alterations in ABC-enriched CD21⁻CD23⁻ DN B cells under physiological aging conditions.

**Conclusions:** Human ABCs appear to comprise a heterogeneous constellation of various subsets possessing different functional properties, as well as different pro-inflammatory or immunoregulatory statuses. ABCs exhibit a distinct hypermetabolic state associated with lupus-related immune dysregulation, but also with aging, in line with previous reports. However, as the disease manifests and progresses, protein production appears to be dampened, which could be attributed to ribosomal problems or proteasome-mediated protein degradation. Metabolic reprogramming is accompanied by enhanced phagocytic and lysosomal programs, which may facilitate the uptake of self-material and contribute to autoreactive immune responses, in particular anti-nuclear autoantibody production. Increased OXPHOS activity may accommodate epigenetic reprogramming towards the atypical memory phenotype displayed by ABCs. Identification of ABC-specific metabolic dependencies may provide opportunities for therapeutic intervention, with lipid consumption being of particular interest. These findings support further investigation of metabolic modulators, including HMG-CoA reductase inhibitors and proteasome-targeting strategies, to limit their pathogenic activity in SLE by exploiting their niche metabolic vulnerabilities, in particular their hypermetabolic state and proteostatic stress.

## 1. Introduction

Age-associated B cells (ABCs) are a heterogeneous population of antigen-experienced memory B cells that accumulate during normal aging, but also expand significantly in autoimmune diseases. Rather than representing a single fixed lineage, ABCs are increasingly viewed as a type of inflammation-induced memory B cell, generated by chronic antigen exposure (BCR), innate stimulation (TLR7 and 9), and inflammatory cytokines (IFNγ, IL-21) **(Rubtsov et al., 2015; Naradikian et al., 2016)**, and have emerged as a niche target of interest in various autoimmune pathologies **(Cancro, 2020; Mouat et al., 2022).** ABCs were initially defined as mature, non-marginal zone and non-follicular B2 B cells that lack CD21, CD23, CD95, and CD43 **(Hao et al., 2011)**, and were later shown to express the integrins CD11b and CD11c, as well as the transcription factor T-bet, in parallel with the upregulation of T cell costimulatory and antigen-presenting molecules, such as CD80, CD86, and MHCII **(Rubtsov et al., 2011; Rubtsova et al., 2015)**. ABCs are commonly identified as CD19^+^B220^+^CD21^-^CD23^-^, with increased expression of the CD11b and CD11c integrins **(Rubtsova et al., 2015; Hassani et al., 2026)**. Besides T-bet, Zeb2 was recently described as another signature transcription factor that shapes the ABC phenotype, additionally endowing them additionally with unique functional properties, such as enhanced viability, migration potential, and phagocytosis/engulfment **(Dai et al., 2024)**. In humans, the corresponding equivalent population can be found among the atypical memory fraction of IgD^-^ CD27^-^ B cells **(Beckers et al., 2023)**.

In autoimmune settings, a large body of literature implicates ABCs in the pathogenesis of systemic lupus erythematosus (SLE) **(Hassani et al., 2026; Z. Li et al., 2023; Mouat et al., 2022; Nickerson et al., 2023)**. SLE is a chronic autoimmune disease in which maladaptive autoimmune responses are directed against multiple organs, resulting in tissue damage, affecting primarily women **(Kaul et al., 2016).** SLE’s complex and multifactorial pathogenesis involves dysregulated interactions among several immune-cell populations, including B cells, neutrophils, and dendritic cells, which promote autoreactivity through the production of autoantibodies and pro-inflammatory mediators **(Tsokos, 2024).** Despite advancements in treatment options that have contributed to tremendous progress in the treatment of SLE, cases of refractory or relapsing SLE necessitate the discovery of alternative pharmacotherapeutic avenues. ABCs have emerged as potential targets for SLE treatment and, moreover, manipulation of immunometabolic properties of SLE-driving immune cells, in particular B cells, is emerging as a promising toolbox to complement SLE treatment **(Bettencourt & Powell, 2017; Makowski et al., 2020; Ueno et al., 2026)**

Metabolic reprogramming is instrumental in immune cell activation, ensuring the provision of adequate ATP and building-block metabolites for immune cells to perform their immune-effector functions. Glucose uptake is essential for B cell adaptive responses, since B cell–specific knock-out of the glucose transporter GLUT1 reduced B cell numbers and impaired immunoglobulin production *in vivo* **(Caro-Maldonado et al., 2014),** in parallel with reduced generation of germinal center B cells and antibody-secreting cells **(Bierling et al., 2024)**. Additional murine studies have monitored metabolic changes across differentiation lines, following activation of naive B cells, wherein B cell activation led to upregulation of glycolysis, followed by an incremental increase of OXPHOS during plasma cell differentiation **(Bradford et al., 2024; Lam et al., 2016; Price et al., 2018)**.

Various studies have previously reported profound metabolic reprogramming taking place across diverse immune cell populations in SLE, including T cells, germinal center B cells, and myeloid cells. These alterations encompass enhanced glycolysis and oxidative phosphorylation, dysregulated amino acid and lipid metabolism, aberrant PI3K-AKT-mTOR signalling, and impaired redox homeostasis. Mitochondrial dysfunction and the subsequent accumulation of reactive oxygen species further amplify inflammatory responses and promote tissue damage, thereby contributing to SLE pathogenesis **(Patiño-Martinez & Kaplan, 2025).** BAFF, a known driver and therapeutic target in SLE, propagates signals through the metabolic PKCβ and AKT pathways and expands metabolic capacity and glucose metabolism in chronically triggered B cells **(Caro-Maldonado et al., 2014; Patke et al., 2006; Wallace et al., 2009),** while inhibition with rapamycin (mTOR inhibitor) can inhibit BAFF-stimulated B-cell proliferation and survival **(Zeng et al., 2015)**. Another study framed the monocarboxylate transporter 1 (MCT1) as a potential target to dampen hypermetabolic activity in SLE, since MCT1 serves as a metabolic gatekeeper for metabolic fluxes of lactate and pyruvate across the plasma membrane, which then enables class-switch recombination to IgG subtypes, through an epigenetic mechanism that involves H3K27 acetylation, hence establishing a connection between metabolic reprogramming and B cell humoral responses **(Chi et al., 2024; Patiño-Martinez & Kaplan, 2025)**. Thus, pharmacological modulation of selected dysregulated metabolic pathways, including glycolysis, glutamine metabolism, fatty acid synthesis or oxidation, the tricarboxylic acid cycle (Krebs cycle), the mitochondrial electron transport chain, could represent a promising strategy for treating autoimmune diseases **(Bettencourt & Powell, 2017; Bradford et al., 2024)**.

In SLE, B cells are chronically exposed to signals mediated by the B cell receptor, Toll-like receptors, interferons, and other cytokines, which can potentiate and alter glycolysis, mitochondrial respiration, lipid metabolism, redox balance, and protein synthesis **(Berry et al., 2020; Lauterbach et al., 2019; D. Wu et al., 2016)**. Additional SLE studies, utilizing the triple congenic (TC) SLE murine model, have also shown that ABC display a hypermetabolic phenotype in parallel with ROS production, while treatment with metformin, a prominent antidiabetic metabolic modulator, was able to suppress this metabolic hyperactivity and achieve disease remission **(Ramirez De Oleo et al., 2023)**.

Regarding ABCs, elucidating the immunometabolic properties of ABCs in particular is an expanding area of research. Recent aging studies have reported a hypermetabolic phenotype in ABCs, identified as mature CD21⁻CD23⁻ double-negative (DN) B cells, characterized by increased glucose uptake, in addition to enhanced oxygen consumption rate (OCR) and extracellular acidification rate (ECAR) measurements, as measured using the SEAHORSE technology **(Frasca et al., 2021)**. The same group also showed that obesity-driven metabolic inflammation, a condition characterized by various metabolic perturbations, such as hyperglycaemia and hyperlipidaemia, can drive production of autoantibodies. In vitro incubation of B cells from young, lean individuals with the fatty acid palmitate induced transcription of T-bet and triggered the secretion of IgGs against adipocyte-derived antigens, such as malondialdehyde **(Frasca et al., 2022)**, suggesting an interconnection between lipid substrates and the phenotype of ABCs.

In the present study, we combined comprehensive transcriptomic analyses with functional metabolic profiling to characterize the immunometabolic programs of ABCs in aging and SLE, two major contexts associated with their expansion. Re-analysis of scRNA-sequencing datasets of peripheral blood mononuclear cells (PBMCs) from SLE patients revealed substantial intra-ABC heterogeneity and identified three distinct non-canonical subpopulations characterized by the expression of NKG7, CD68, or Areg. Despite B cells displaying a hypermetabolic phenotype, ex vivo metabolic profiling with SCENITH showed dampened global protein synthesis, which may serve as a point for therapeutic intervention through drug repurposing of metabolic modulators to target the ribo-proteinogenic infrastructure of pathogenic B cells. Further transcriptomic analysis suggested increased proteasome degrading activity may account for the lower protein synthesis capacity, observed in B cells from active SLE. Finally, high-resolution metabolic analysis of B cells from cross-examining aging mice identified altered NADH- and FAD-associated redox states accompanying ABC expansion, linking age-dependent changes in cellular redox metabolism to the emergence of this population.

## 2. Materials and Methods

### 2.1 Public scRNA-seq datasets retrieved from the Gene Expression Omnibus (GEO) repository

For transcriptomic re-analysis of age-associated B cells (ABCs) in the human condition, we retrieved the dataset GSE163121, generated by **(Bhamidipati et al., 2021).** This dataset was integrated with the GSE242615 dataset, which was generated by the groups of Vinuesa & Shen **(Dai et al., 2024).** To examine deeper ABCs in the context of the murine SLE-like MRL.lpr model, we used the dataset GSE202110, generated by the group of Mark Shlomchik **(Nickerson et al., 2023).**

### 2.2 Computational pipelines for scRNA-seq re-analysis

#### 2.2.1 Packages and software environment

Raw single-cell RNA-sequencing data generated by the 10x Genomics platform were processed using Seurat for pre-processing, clustering, and cell-state identification **(Hao et al., 2021).** ScRNA-seq analysis was conducted on Seurat v3. Potential technical artifacts/doublets were removed using DoubletFinder **(McGinnis et al., 2019)**. Differential expression, pathway enrichment, disease ontology analysis, and gene set enrichment were performed using fgsea **(Korotkevich et al., 2016)**, clusterProfiler **(Yu et al., 2012),** pathviewR **(Baliga V & Armstrong M, 2021)** and DOSE **(Yu et al., 2015)**. Intercellular communication networks were inferred using CellChat **(Jin et al., 2025)**, while cellular state transitions and lineage pseudo-temporal relationships were reconstructed with monocle3 **(Cao et al., 2019)**. Gene co-expression programs were identified using hdWGCNA **(Morabito et al., 2023)**, and regulatory and pathway activities were inferred with decoupleR **(Badia-i-Mompel et al., 2021)**. Metabolic-specific pathway analysis was investigated using scMetabolism **(Y. Wu et al., 2022)**. Drug repurposing candidates and their target networks were identified using DMEA, SCAnet, and ASGARD **(Garana et al., 2023; He et al., 2023; Oubounyt et al., 2023)**. Metabolite-mediated intercellular communication was analysed in Python using MEBOCOST **(R. Zheng et al., 2025)**. Visualization was enabled by the ggplot2 package **(Wickham, 2016)**.

#### 2.2.2 Workflow for scRNA-seq analysis

Single-cell RNA sequencing datasets from two healthy controls (HC1 and HC2) and four systemic lupus erythematosus (SLE) samples (SLE1–SLE4) were imported using the Read10X function. Individual Seurat objects were generated for each sample using the CreateSeuratObject function with filtering thresholds of a minimum of three cells per gene and at least 200 detected genes per cell. Metadata identifying disease status (healthy control or SLE) were added to each object. All samples were merged into a single Seurat object for downstream analyses. Cells were retained if they met the following criteria: 200 < detected genes < 5,000, Total UMI counts < 10,000, mitochondrial transcript percentage < 12% and haemoglobin transcript percentage < 0.0001. Gene expression values were normalized using Seurat’s NormalizeData function. Highly variable genes were identified using the variance-stabilizing transformation (VST) method implemented in FindVariableFeatures, with the top 4,000 variable features retained for downstream analyses. Cell cycle effects were evaluated using canonical S-phase and G2/M-phase gene sets provided by Seurat. CellCycleScoring was applied to assign each cell a cell-cycle phase and corresponding scores. Gene expression values were scaled using ScaleData while regressing out unwanted sources of variation, including: Mitochondrial transcript percentage, Ribosomal transcript percentage, Immunoglobulin transcript percentage. Principal component analysis (PCA) was performed on the scaled expression matrix. The optimal number of principal components was evaluated using elbow plots and heatmaps of principal component loadings. A shared nearest-neighbor graph was constructed using the first 45 principal components. Cells were clustered using the Louvain algorithm across multiple resolution parameters (0.2–2.0), and cluster stability was assessed using the *clustree* package **(Zappia & Oshlack, 2018)**. A final clustering resolution of 1.2 was selected. Uniform Manifold Approximation and Projection (UMAP) was applied using the first 45 principal components to visualize cellular heterogeneity in two-dimensional space. Potential doublets were identified using DoubletFinder. Parameter optimization was performed using the paramSweep and find.pK functions to identify the optimal pK value. The expected doublet rate was estimated as 10% of total cells and adjusted for homotypic doublets using the modelHomotypic function. Doublets were classified using DoubletFinder with the following parameters: (Principal components: 1–45, pN = 0.30, pK = 0.01). Following doublet removal, datasets were split by sample identity. Each dataset was independently normalized and variable genes were identified. Integration features were selected using SelectIntegrationFeatures, and integration anchors were identified using canonical correlation analysis (CCA) through FindIntegrationAnchors. Datasets were integrated using IntegrateData to correct for inter-sample batch effects. The integrated dataset was subsequently scaled, subjected to PCA, and visualized using UMAP. Quality control checks can be found in **(Supplementary Figure S1)**.

#### 2.2.3 Developmental cell fate trajectory analysis with monocle3

Trajectory analysis was performed using the Monocle3 R package **(Cao et al., 2019)**. Gene expression dynamics along pseudotime were visualized using the plot_genes_in_pseudotime() function from the Monocle3 package. Analysis was restricted to selected B-cell populations (early B cells, B cells, ABCs, and plasma cells). Genes of interest were plotted across the pseudotime differentiation line, and cells with expression levels below 0.1 were excluded from visualization (min_expr = 0.1). Expression trends across pseudotime were modelled using a natural cubic spline with 7 degrees of freedom *(trend_formula = "∼ splines::ns(pseudotime, df = 7)")*, allowing flexible fitting of non-linear gene expression patterns during B-cell differentiation.

#### 2.2.4 Cell-cell communication analysis with CellChat

Cell–cell communication analysis was performed using the CellChat R package. Cells from the SLE group were extracted from the integrated Seurat object The CellChat human ligand–receptor database (CellChatDB.human) was used, including *“Secreted Signalling, ECM–Receptor, and Cell–Cell Contact”*. Overexpressed signalling genes and ligand–receptor pairs were identified using CellChat’s default workflow. Intercellular communication probabilities were inferred from normalized gene expression profiles using the triMean method implemented in computeCommunProb(). Cell populations containing fewer than three cells were excluded from downstream analyses (min.cells = 3).

#### 2.2.5 Metabolite-driven cell-cell communication with MEBOCOST

Metabolite-mediated cell–cell communication was inferred from single-cell RNA-sequencing data using MEBOCOST. A pre-processed PBMC dataset containing cells from systemic lupus erythematosus (SLE) patients and healthy control individuals (HC) was analysed with cell-type annotations and condition labels supplied as metadata. Communication events were inferred based on the coordinated expression of metabolite-producing enzymes in sender cells and metabolite sensors in receiver cells. Statistical significance was evaluated using 1,000 permutations, and significant interactions were identified using permutation-based FDR correction. Differential communication analysis was performed between SLE and HC to identify disease-associated changes in metabolite signalling. Results were visualized using network, flow, scatter, and summary plots to characterize alterations in intercellular metabolic communication across immune cell populations.

#### 2.2.6 KEGG pathway analysis with *Clusterprofiler* and *scMetabolism*

Differentially expressed genes were identified using Seurat’s FindMarkers function. Gene identifiers (SYMBOL IDs) were converted to Entrez IDs using the clusterProfiler bitr function, and unmapped or duplicated entries were removed. A ranked gene list was constructed using log2 fold-change values and used for KEGG gene set enrichment analysis with the gseKEGG function (clusterProfiler), applying 10,000 permutations, a gene set size range of 3–1200 genes, and organism-specific annotation (hsa), with Entrez Gene IDs as the key type. KEGG pathway visualization was performed using the pathview R package. For more specialized metabolic pathway analysis, the scMetabolism package was used with default settings. Briefly, metabolic pathway activity scores were computed from the UMI count matrix using the *sc.metabolism.Seurat* function with the VISION algorithm. No imputation was applied, and analyses were performed using a KEGG-defined metabolic gene set gmt.file (85 pathways). Pathway activity scores were extracted from the METABOLISM assay slot of the Seurat object. Visualization of metabolic signatures was performed using scMetabolism built-in functions for dimensional reduction and pathway-level plotting.

#### 2.2.7 Metabolic pathway scoring and binary plotting

To contextualize metabolic pathway dynamics, we plotted metabolic pathways in relevant binary combinations, by adapting an approach previously described in **(Xiao et al., 2019)**. In brief, metabolic and signalling pathway activity scores were computed from single-cell RNA-seq data by applying gene set variation analysis based on predefined KEGG-derived metabolic pathways and MSigDB Hallmark gene sets obtained using gmtPathways. Gene sets (e.g., glycolysis, oxidative phosphorylation, TCA cycle, lipid pathways) were intersected with genes detected in the Seurat object to ensure compatibility with the expression matrix, using the *intersect()* function. Normalized expression data were extracted from the Seurat assay (GetAssayData, layer = "data"), and pathway scores were calculated at single-cell resolution as the mean expression of all genes within each signature using *Matrix::colMeans* function. These scores were integrated with cell identity annotations and sample metadata to enable group-wise comparisons, using the *data.frame()* function. Relationships between metabolic programs were visualized using scatter and/or density-based plots.

### 2.3 Animal models

In the aging experiments, young (<12 weeks old) and old (> 80 weeks old) wild-type male mice were used on the C57BL/6J background. Those mice were bred in the TARC (Translational Animal Research Center of the University Medical Center Mainz) facility under specific pathogen-free (SPF) conditions. For the lupus MRL.lpr experiments, tissue samples were kindly provided by the group of Prof. Dr. med Julia Weinmann-Menke, in line with the 3R principles. All animal experiments were performed in accordance with the institutional regulations of the Central Animal Facility.

### 2.4 Preparation of single cell suspension for flow cytometry

To prepare a single cell suspension, spleens of MRL.lpr mice were smashed through a 40μm filter in flow cytometry staining buffer (PBS buffer supplemented with 2% fetal calf serum). Cells were centrifuged at 300g for 7 minutes and resuspended in 1mL od ACK lysis buffer to lyse erythrocytes for approximately 3 minutes. Lysis reaction was terminated by adding 10mL of staining buffer. First, Fc-block (5 µg/mL) was used for 15 min at 4 °C, followed by surface marker staining, for 20 min at 4 °C with antibodies against: CD23-PE (Biolegend: #101608), CD45R/B220-Percp (Biolegend: #103234), CD19-PECy7 (Biolegend: #115520), Tbet-efl660 (eBioscience: #46-0333), CD21/35-PacificBlue (Biolegend: #123414), VD EFL506 (eBioscience: #65-0866), CD5 – BUV737 (BD Biosciences: #612809), CD45R/B220-BV785 (BD Biosciences: #103246), CD90.2-RB780 (BD Biosciences: #755824), CD23-PECy7 (Biolegend: #115520), CD19-APCR700 (BD Biosciences: #565473), CD21-APCCy7 (BioLegend: #123418).

### 2.5 Metabolic analysis with the SCENITH assay

We implemented the SCENITH (Single-Cell Energetic Metabolism by Profiling Translation Inhibition) assay as previously described **(Argüello et al., 2020)**. In brief, cells from each male and female MRL.lpr sample were cultured in quadruplicates in full DMEM medium in a 96-well U plate. Cells were incubated in the presence of 2DG (100mM), Oligomycin (2μM), 2DG+Oligomycin (100mM, 2μM) or DMEM medium only (Baseline steady-state), for 20-30 minutes in a 37°C CO2 incubator. The puromycin solution was added at 2x for a final concentration of 10μg/mL, for another 30min to allow for puromycin incorporation in newly synthesised proteins. The plate was centrifuged for 7min at 300g, cells washed with FACS buffer, and the antibody staining protocol was followed. Cells were fixated and permeabilized with the FOXP3 kit and stained intracellularly with an Alexa647-conjugated anti-puromycin antibody (Sigma-Aldrich) for 60 minutes, following the manufacturer instructions. After incubation, cells were washed with the kit’s permeabilization buffer (1x), followed by resuspension in FACS acquisition buffer. Samples were acquired with a FACS Canto II flow cytometry instrument (BD Bioscience).

### 2.6 Redox balance analysis of NADH and FAD

#### 2.6.1 Sample preparation for redox balance analysis

Approximately 3-5x10^6^ spleen cells were seeded on a 96-well U-plate from young and old mice. Following centrifugation at 300g for 6 minutes, the Fc receptors were blocked with anti-mouse CD16/32 (BioLegend) for 15 min at 4 °C in the dark. Following washing with FACS buffer, cells were stained with an antibody cocktail in FACS buffer (Table S1) for 20-30 minutes at 4 °C in the dark. Cells were then washed twice in FCS-free DMEM medium (Thermo Fisher), centrifuged, and resuspended in glucose-containing FCS-free DMEM medium at a final volume of 500 µL in the presence of viability dye 7-AAD (BD Bioscience) to exclude cells dying through the mitostress test, and were left to equilibrate at room temperature to prevent temperature-induced metabolic shifts.

#### 2.6.2 Sample acquisition and NADH/FAD autofluorescence measurement

Redox balance analysis was conducted as previously described **(Bartolomé & Abramov, 2015; Abir et al., 2024; Lemire et al., 2022; Stylianakis & Hövelmeyer, 2026)**. In brief, bulk splenocytes were acquired at baseline, followed by sequential additions of oligomycin (20uM), FCCP (10uM), and rotenone/antimycin A (2uM) at approximately 30-second intervals with short, gentle vortex. Samples were analysed with a FACS Symphony A5 spectral unmixer (BD Bioscience). FCS files were analysed using the FlowJo v10.10 software. The antibody panels were designed as such to minimize spill-over into the channels used to measure NADH and FAD autofluorescence excited by the 355-nm UV laser and 488-nm blue laser, respectively **(Stylianakis & Hövelmeyer, 2026)**. To augment autofluorescence intensity, detector voltages were incrementally increased to ensure that signals remained within the positive range. To circumvent spill-over issues of NADH to the FAD signal from UV excitation, FAD was excited with a 488-nm blue laser, which does not excite NADH. The ORR was calculated as [FAD/(NADH+FAD)], To further estimate absolute metabolic activity per cell, the side scatter parameter was factored into ORR, based on the positive correlation between mitochondrial mass and granularity.

### 2.7 Measurement of mitochondrial mass and mitochondrial membrane potential (MMP)

Spleens were harvested from young and old C57BL/6 mice, and a single-cell suspension was prepared as described in the method section above. Following surface staining and incubation in a 96-well U plate (Thermo Fischer), cells were washed with PBS^+/+^ Ca^2+^/Mg^2+^ to remove traces off fetal calf serum, and were stained with MiotrackerGreen (Thermo fisher, dil.ratio 1:5000) and tetramethylrhodamine ethyl ester perchlorate (TMRE; Abcam, dil.ratio 1:5000) in non-quenching mode, to measure mitochondrial mass and mitochondrial membrane potential respectively, following established practises **(Connolly et al., 2018)**. Cells were incubated in PBS^+/+^ (Ca^2+^/Mg^2+^) buffer (Thermo Fisher), in the presence of 7-AAD to exclude dead cells, in the dark at room temperature for 20 minutes. Following incubation, cells were washed twice with staining buffer. Samples were acquired immediately upon resuspension in staining buffer, in a FACSymphony A5 SE flow cytometry instrument.

### 2.8 Imaging flow cytometry for visualization of MitotrackerGreen, TMRE and granularity

Spleen cells from young and old mice were stained with MitoTracker Green FM, TMRE, and Hoechst 33342 according to the manufacturers’ recommended conditions, to visualize mitochondrial parameters in CD21^-^CD23^-^ CD19^+^ B cells with all staining and subsequent handling performed under light-protected conditions. Following staining, cells were washed with and resuspended in staining buffer at an appropriate concentration for acquisition. Samples were analyzed using an ImageStreamX Mk II imaging flow cytometer (Cytek) at 60X magnification. Acquired raw imaging (rif) files were analyzed using the IDEAS software, with focused, single cells identified based on image quality and morphology, followed by gating of Hoechst-positive cells. Side-scatter area (SSC-A) was also acquired and visualized as a measure of cellular granularity and internal complexity.

### 2.9 Statistical analysis

All experiments undergoing statistical analysis included at least three or more biological replicates. Statistical analyses and figure generation were performed with Prism v10 software (GraphPad). Statistical significance between groups was calculated with two-way ANOVA with Šidák’s multiple comparisons analysis, or Student’s t test (paired/unpaired) where applicable. Statistical tests were applied as indicated in the figure legends. All values are represented as mean ± SEM unless otherwise stated. P values were considered significant with p<0.05. No blinding of mice or samples was conducted. At least 3 mice were used per experiment. No statistical methods were used to determine the sample size.

## 3. Results

### 3.1 Single-cell transcriptomic profiling resolves age-associated B cell heterogeneity in SLE patients

To precisely characterize how cell-intrinsic metabolic reprogramming dictates the distinct functional states of atypical memory age-associated B cells (ABCs) during chronic autoimmunity, we performed a re-analysis of a scRNA-seq dataset corresponding to sorted peripheral blood mononuclear cells-derived B cells from systemic lupus erythematosus (SLE) patients and healthy control (HCs) individuals **(Fig.1A).** Unsupervised clustering and Uniform Manifold Approximation and Projection (UMAP) identified major immune cell lineages based on canonical marker expression, including early B cells, mature B cells, ABCs, and plasma cells, alongside minor populations of T cells, NKT cells, CD8^+^ T cells, and monocytes/macrophages (**Fig. 1A).** Comparative analysis between the experimental groups revealed striking shifts in the cellular composition and population density of the B-cell compartment. As shown in the original studies **(Bhamidipati et al., 2021; Q. Liu et al., 2023),** in healthy control individuals (HC), ABCs constituted a distinct, restricted cluster segregated from the primary pool of naive/mature B cells (**Fig. 1A, left panel**), while profiling of SLE patient-derived PBMCs demonstrated a profound expansion of the ABC cluster (**Fig. 1A, right panel, and 1B**) **(Bhamidipati et al., 2021; Q. Liu et al., 2023)**. This accumulation was accompanied by a prominent increase in plasma cell density, mirroring active differentiation and sustained disease escalation **(Dang et al., 2025)**. Further sub-clustering of the ABC cluster revealed two subpopulations, termed 0 (red) and 1 (green), that expanded and were present only in the SLE group, as shown in the UMAP illustration **(Fig. 1B)**.

**Figure 1.**
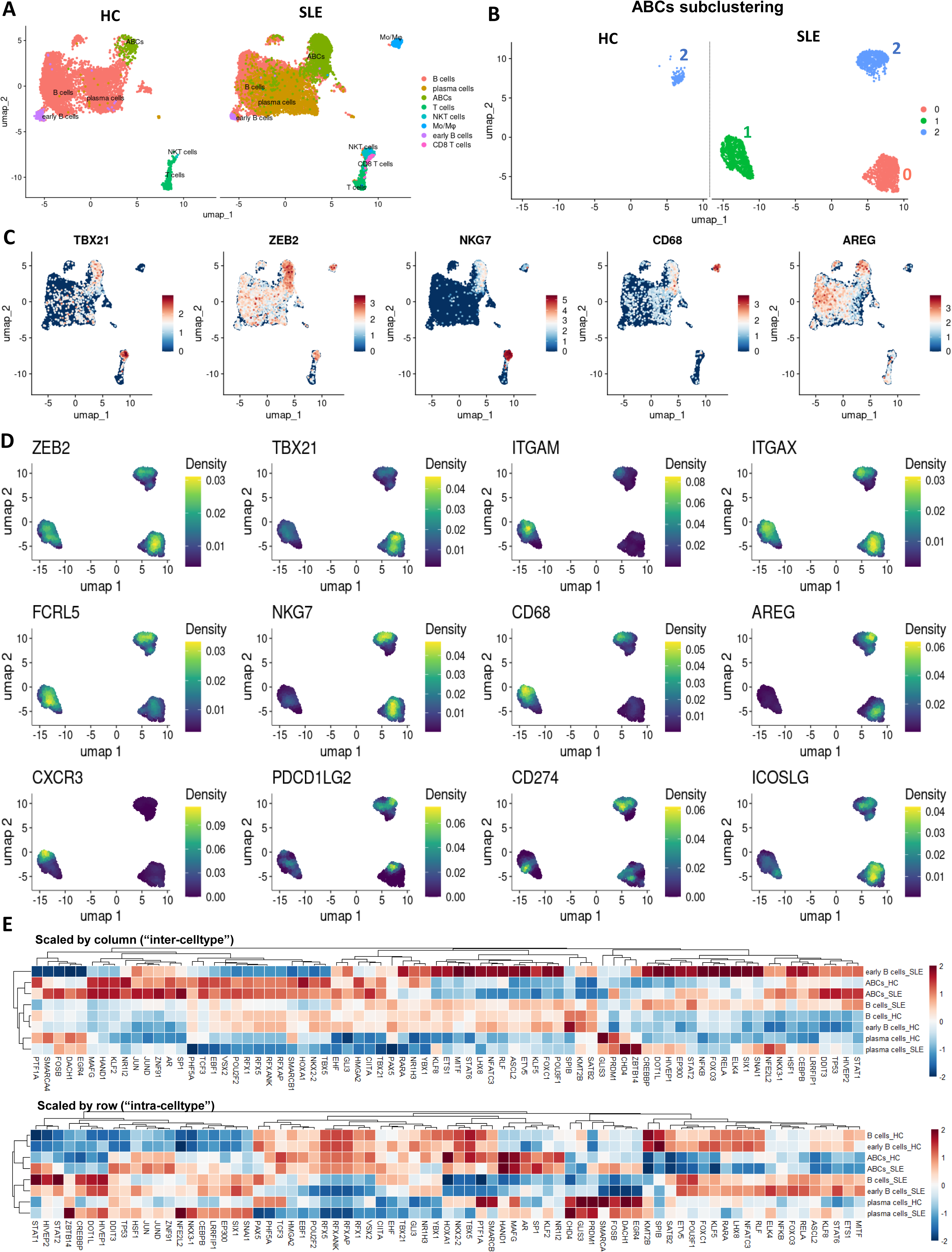
scRNA-seq reanalysis of PBMC-derived B cells from two healthy donors and four patients with SLE. **(A–B)** UMAP visualization of immune cell populations and ABC subclusters between healthy controls (HC) and SLE individuals (SLE). **(C)** Immunophenotypic annotation and characterization of the ABC cluster, with visualization of TBX21, ZEB2, NKG7, CD68, AREG. **(D)** Immunophenotypic characterization of the three ABC subclusters. **(E)** Transcription factor activity profiling inferred using the *decoupleR* package. ***Upper panel:*** column-scaled values (intercellular comparison). ***Lower panel:*** row-scaled values (intracellular comparison).

The ABC cluster was annotated by the co-presence of the well-established signature transcription factors TBX21 (T-bet) and ZEB2 **(Fig. 1C) (Dai et al., 2024; Hassani et al., 2026; Z. Li et al., 2023)**, wherein we also observed co-expression of NKG7, CD68 and amphiregulin (AREG), molecules commonly associated with natural killer cytotoxicity, phagolysosomal activity, and regulatory or suppressive activity respectively **(Chistiakov et al., 2017; Ehnold et al., 2025; Turiello et al., 2025)**. However, we observed **(Fig. 1D)** profound immunophenotypic variability with regard to ZEB2, T-BET, ITGAM (CD11b), ITGAX (CD11c), FCRL5, as well as NKG7, CD68, and AREG, among the three ABC subclusters. These markers helped differentiate and distinguish among the three ABC subclusters (CellType: 0, 1, and 2), which indicates an underappreciated plasticity within age-associated B cells **(Fig. 1D)**.

We further conducted transcription factor analysis with the decoupleR package, which clearly showed that the ABC cluster possesses a unique and distinctive transcription factor fingerprint, relative to the other B cell populations **(Fig. 1E)**. Hierarchical clustering of transcription factor expression, both across cell types (**Fig. 1E, upper panel)** and within each cell type (**Fig. 1E, lower panel)**, demonstrates that B-cell subsets segregate primarily according to cell identity, with ABCs forming a distinct transcriptional cluster independent of disease status. ABCs are characterized by increased expression of the canonical ABC regulator TBX21 (T-bet) together with ZEB2, ZBTB32, BATF, ID2, and the PRDM1 (Blimp1) plasma cell commitment factor, while exhibiting reduced expression of core B-cell lineage factors such as PAX5 **(Fig. 1E)**.

### 3.2 Identification of gene regulatory networks in lupus ABCs, associated with cell interaction, metabolic activity and epigenetic reprogramming

To gain a deeper understanding of the gene regulatory circuits that govern ABC behaviour and functionality, we performed high-dimensional weighted gene co-expression network analysis using the hdWGCNA R package, which identifies and constructs context-specific co-expression networks and detects transcriptional modules across cellular clusters **(Morabito et al., 2023)**. After tuning the program with the appropriate parameters, hdWGCNA analysis yielded 3 modules representative of the ABC cluster (ABCs-M1, ABCs-M2, ABCs-M3) **(Fig. 2A-B)**. Downstream regulatory network analysis identified predicted targets that are potentially regulated by TBX21 **(Fig. 2C)** and STAT3 **(Fig. 2D)**. Together, these findings support a model in which ABCs adopt an activated, inflammatory state through the combined action of T-bet- and STAT3-dependent regulatory networks. TBX21 is a well-established regulator of ABC identity that integrates IFN-γ/TLR-driven type 1 responses **(Naradikian et al., 2016)**, whereas STAT3 links cytokine signalling (e.g., IL-6) to STAT3-downstream transcriptional activation.

**Figure 2.**
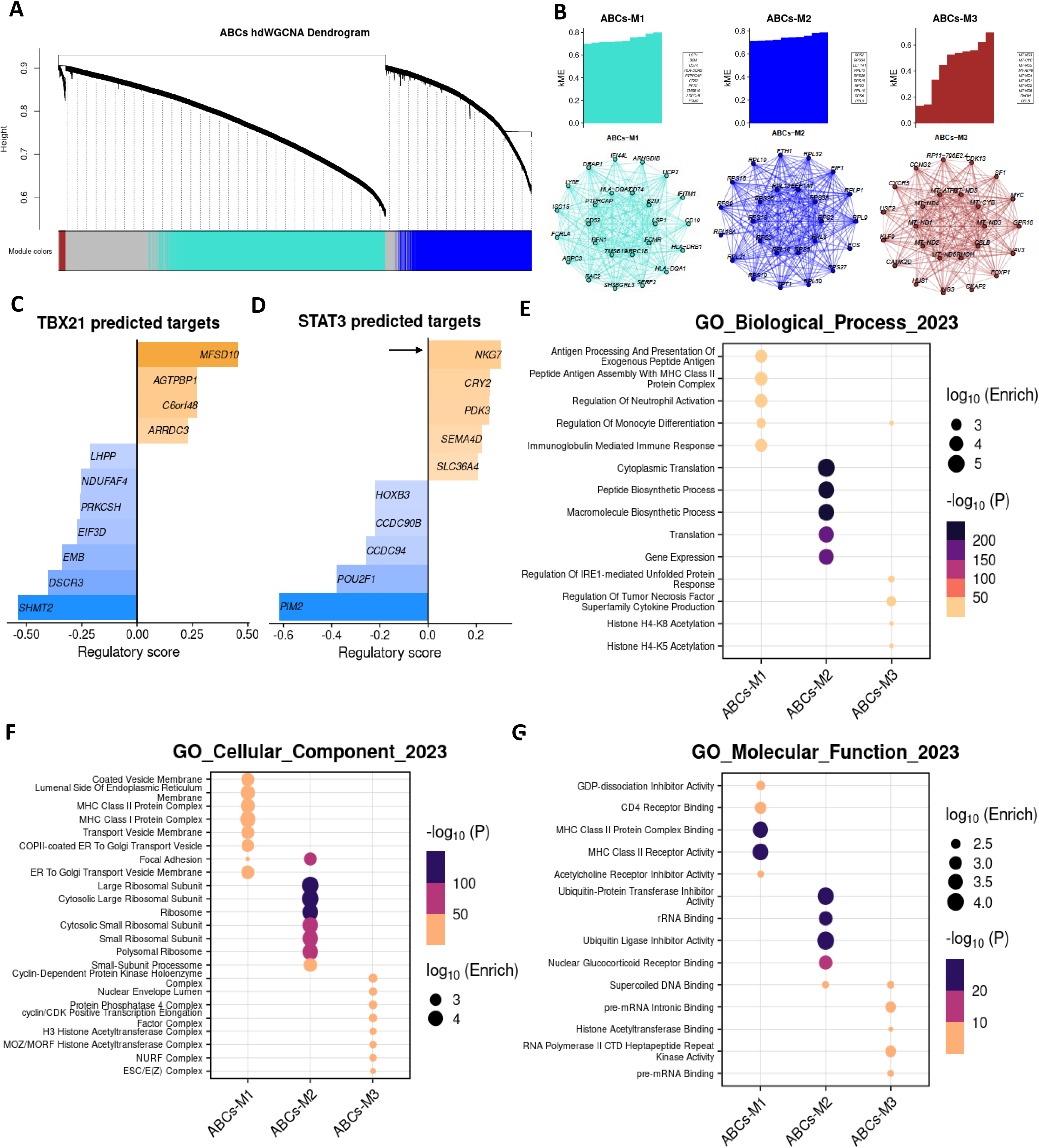
High-dimensional weighted gene co-expression network analysis (hdWGCNA) of age-associated B cells (ABCs). **(A)** Hierarchical clustering dendrogram showing the three transcriptional co-expression modules identified in ABCs (ABCs-M1, ABCs-M2, and ABCs-M3). **(B)** Individual module network bar and radial plots illustrating representative signature genes for the three transcriptional modules (ABCs-M1, ABCs-M2, and ABCs-M3)**. (C–D)** Transcription factor regulatory network analysis centered on TBX21 and STAT3. **(E–G)** Dot plot visualization of Gene Ontology enrichment analysis for each transcriptional module, including biological process (BP), cellular component (CC), and molecular function (MF).

Predicted target genes of TBX21 and STAT3 were ranked by regulatory score, with positive scores indicating predicted activation and negative scores indicating predicted repression. The highest-scoring activated and repressed targets for each transcription factor are shown **(Fig 2C-D)**. T-bet was found to regulate various metabolic aspects. In particular, T-bet appeared to be positively associated with MFSD10/*SLC75A1* (Major Facilitator Superfamily Domain Containing 10), AGTPBP1 (ATP/GTP Binding Carboxypeptidase 1) and ARRDC3 (Arrestin Domain Containing 3), while exhibiting negative correlations with genes involved in mitochondrial function, protein translation, and cellular metabolism, including NDUFAF4 (NADH:Ubiquinone Oxidoreductase Complex Assembly Factor 4), EIF3D (Eukaryotic Translation Initiation Factor 3 Subunit D), and SHMT2 (Serine Hydroxymethyltransferase 2) **(Fig. 2C)**.

STAT3 appears to regulate the expression of Natural Killer Cell Granule Protein 7 (NKG7) **(Fig. 2D)**, which is present in ABC subclusters No0 and No2 **(Fig. 1C-D)**, in addition to genes involved in immune signalling and metabolic regulation, including SEMA4D and PDK3 Moreover, STAT3 was negatively correlated with HOXB3, POU2F1, and PIM2, indicating suppression of alternative transcriptional programs **(Fig. 2D)**. Collectively, these results suggest that T-bet and STAT3 may orchestrate distinct but complementary regulatory networks underlying ABC-cell functional heterogeneity.

Next, we aimed to add further biological context about the performative roles for our co-expression modules by performing different enrichment tests to visualize the hdWGCNA results for the ABC cluster. For the enrichment tests, we utilized the three standard Gene Ontology repositories concomitantly: *<u>GO_Biological_Process_2023</u>* **(Fig. 2E)**, *<u>GO_Cellular_Component_2023</u>***(Fig. 2F)**, and *<u>GO_Molecular_Function_2023</u>* **(Fig. 2G)**. For the ABC-M1 module, the Gene Ontology analysis revealed enrichment of terms related to antigen processing and presentation enabled by MHC-I and MHC-II, cell-cell interaction with monocytes and neutrophils, and molecules associated with focal adhesion (**Fig. 2E-G)**. The ABC-M2 module was composed of terms related to protein translation, ribosomal activity, and ubiquitination, while the ABC-M3 module was reflective of Inositol-requiring enzyme (1IRE1)-mediated unfolded protein response, TNF superfamily cytokine production and epigenetic activity related to histone acetylation (**Fig. 2E-G)**. The most intriguing biological mechanism that was present in the module ABCs-M3, was histone H4-K5/8 acetylation and other epigenetic machinery, such as MOZ/MORF histone acetyltransferase complex, the ATP-dependent nucleosome remodelling NURF complex, and the ESC/E(Z) histone methylating complex. These signatures were shared among healthy individuals and SLE patients alike. Taken together, the ABCs appear to be undergoing active epigenetic rewiring and chromatin architectural remodelling **(Fig. 2E-G)**.

### 3.3 SLE disease is associated with broad-scale metabolic reprogramming in B cells and ABCs

By comparing B cells from the SLE patient group with those from the HC group, we observed a metabolic bimodality and segregation in the utilization of glycolysis and oxidative phosphorylation, respectively, rather than a uniform increase in metabolic activity, as illustrated in the binary pathway scoring plots **(Fig. 3A)**. Regarding lipid metabolism, we found a uniform increase in fatty acid metabolism and cholesterol metabolism in both the ABCs and B cell clusters **(Fig. 3B)**, which, however, was more pronounced among ABCs, suggesting a shift towards lipid consumption or even increased lipid dependence among ABCs in SLE **(Fig. 3C)**. To further characterize the metabolic landscape of B cells and age-associated B cells (ABCs) in patients with lupus, we employed the scMetabolism package for single-cell metabolic pathway analysis **(Y. Wu et al., 2022)**. According to the dot plot illustrating key metabolic pathways across group-stratified B cell subsets **(Fig. 3D)**, ABCs in SLE exhibit a predominantly hypermetabolic phenotype, compared with the rest of B cells, regardless of disease status. This state was characterized by enhanced activity across multiple metabolic pathways, including ketone body metabolism, pyruvate metabolism, the pentose phosphate pathway (PPP), glycolysis, oxidative phosphorylation (OXPHOS), and the tricarboxylic acid (TCA) cycle. In parallel, glycerophospholipid and glycerolipid metabolism, as well as broader cholesterol and fatty acid metabolic programs associated with membrane biosynthesis and lipid raft formation, were markedly upregulated. ABCs also demonstrated increased glutathione metabolism, suggesting mobilization of antioxidant buffers to counteract elevated oxidative stress **(Fig. 3D)**. In contrast, ribosomal rRNA counts were reduced across both conventional B cells and ABCs, as evidenced by decreased expression of ribosomal RNA genes, indicative of a diminished protein synthesis capacity **(Fig. 3E)**. Altogether, ABCs appear to mobilize a wide range of metabolites, nutrients and metabolic pathways to support anabolic and catabolic processes aiming to accommodate not just energy needs but also effector functions (e.g., proliferation, biosynthesis, ROS management).

**Figure 3.**
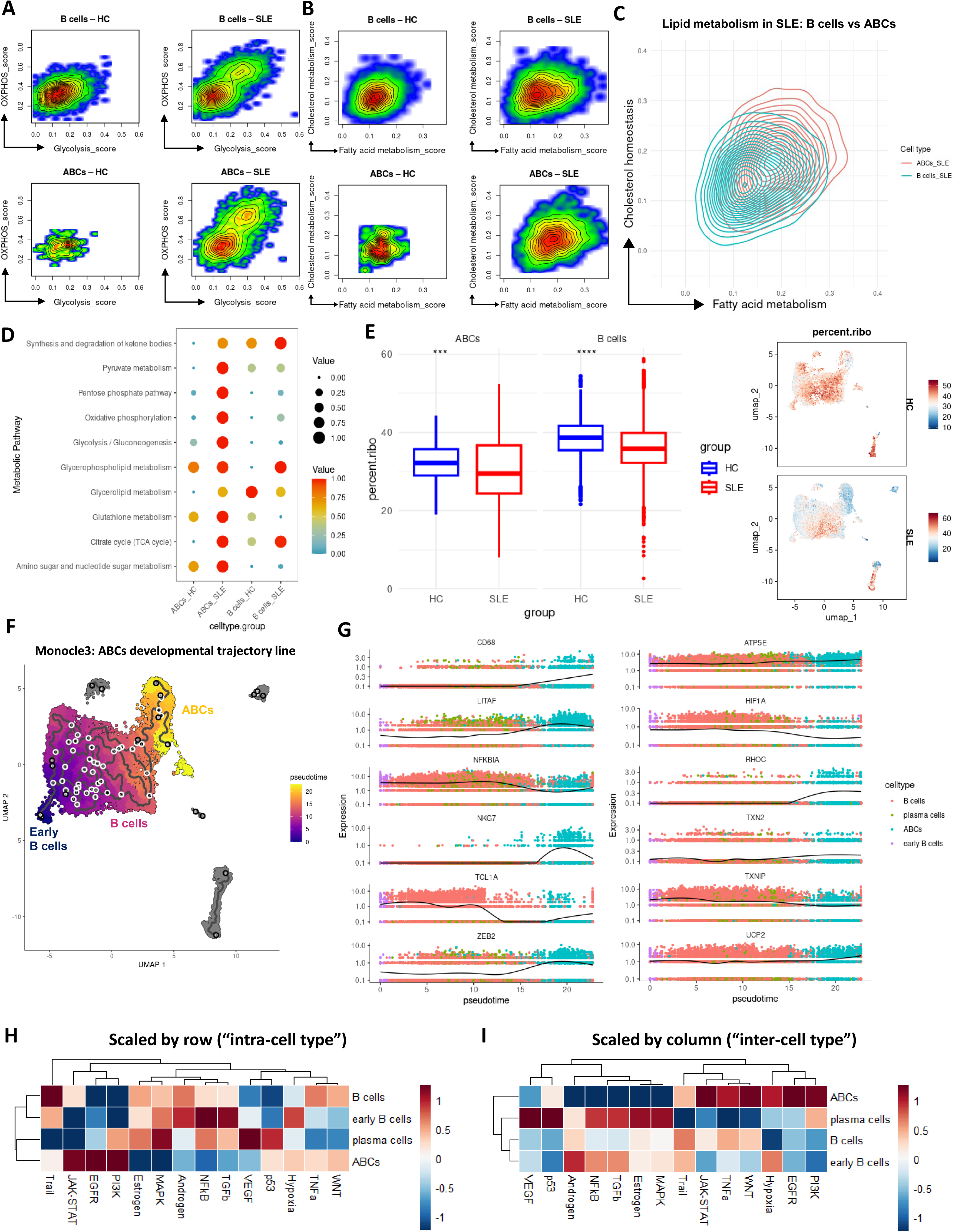
Single-cell RNA sequencing (scRNA-seq)-based metabolic profiling and trajectory analysis of B cells and age-associated B cells (ABCs) from patients with SLE and healthy donors. **(A)** Binary metabolic scoring plots of glycolysis and oxidative phosphorylation in B cells and ABCs from healthy donors and patients with SLE. **(B)** Binary metabolic scoring plots of cholesterol metabolism and fatty acid metabolism in B cells and ABCs from healthy donors and patients with SLE. **(C)** Overlay of binary metabolic scores for lipid metabolism and cholesterol metabolism in SLE-derived ABCs compared with SLE-derived B cells. **(D)** KEGG pathway activity analysis inferred using the *scMetabolism* package. **(E)** Box plot showing ribosomal RNA (rRNA) transcript counts and UMAP visualization of ribosomal transcript expression. **(F)** Monocle3-inferred pseudotime trajectory depicting the differentiation of B cells into ABCs. **(G)** Expression dynamics of selected genes of interest along the pseudotime trajectory. **(H-I)** Transcription factor activity inferred using the **decoupleR** package. **H)** Row-scaled z-scores values (intracellular comparison). **J)** Column-scaled z-scores values

To investigate metabolic reprogramming along differentiation dynamics, we performed pseudotime trajectory analysis using Monocle3, which formed a developmental line starting from early/transitional B cells, crossing mature/activated B cells and effectively culminating in ABCs, at later stages of differentiation **(Fig. 3F)**. Progression along this developmental trajectory was marked by upregulation of the lysosomal marker CD68, the cytotoxic marker NKG7, and the signature transcription factor ZEB2, together with progressive upregulation of the LITAF (Lipopolysaccharide-Induced Tumor Necrosis Factor-Alpha Factor), in parallel with a progressive downregulation of the early B cell factor TCL1A and the inhibitory IkBa (NFKBIA) factor **(Fig. 3G – left panel)**. Interestingly, we also saw upregulation of Rho-kinases (RHOC), which have been postulated as potential therapeutic targets for SLE **(Fig. 3G – right panel) (Q. Liu et al., 2023; Sachinidis et al., 2025; Sachinidis & Garyfallos, 2023; Stirzaker et al., 2012)**. In addition, cell fate decision towards ABCs coincided with increased expression of OXPHOS and electron transport chain–associated genes, including ATP5E and UCP2, together with upregulation of thioredoxin TXN2 and downregulation of the inhibitory Thioredoxin-interacting protein (TXNIP), genes involved in buffering oxidative stress and redox regulation **(Fig. 3G – right panel)**. Pathway analysis with decoupleR showed that ABCs were strongly enriched for proinflammatory and effector pathways, such as JAK-STAT, TNFα, EGFR, WNT, and PI3K signalling pathways, and mildly for p53 and hypoxia, compared to B cells **(Fig. 3K).**

### 3.4 ABC subpopulations exhibit metabolic heterogeneity

To further dissect the metabolic heterogeneity observed within the ABC compartment, we performed binary pathway scoring across the three ABC subclusters (Celltype No0 = NKG7^+^ B cells, Celltype No1 = CD68^+^ ABCs and Celltype No2 = AREG^+^ ABCs) **(Fig. 4A)**. Among these, the CD68^+^ ABC subcluster No1 displayed the most pronounced hypermetabolic phenotype, consistently exhibiting elevated activity across major metabolic pathways, such as glutathione metabolism, the pentose phosphate pathway, lipid and nucleotide metabolism, reactive oxygen species (ROS) generation, TCA cycle activity, glycerolipid and glycerophospholipid metabolism, and extra-mitochondrial pathways such as peroxisomal function and ketone body utilization **(Fig. 4A)**. Marker-based annotation demonstrated that SLE-associated ABCs are heterogeneous and encompass at least three functional states: (i) an inflammatory/activated state <u>(ABCs subcluster 0)</u>, (ii) an interferon-responsive state <u>(ABCs subcluster 1)</u>, and (iii) an antigen-stimulated effector state <u>(ABCs subcluster 2)</u> **(Fig. 4B)**. The co-existence of these transcriptional programs highlights the dynamic nature of ABC differentiation in SLE and aligns with the concept that ABCs integrate both chronic interferon exposure and ongoing antigenic stimulation during disease progression **(Fig. 4B-C)**.

**Figure 4.**
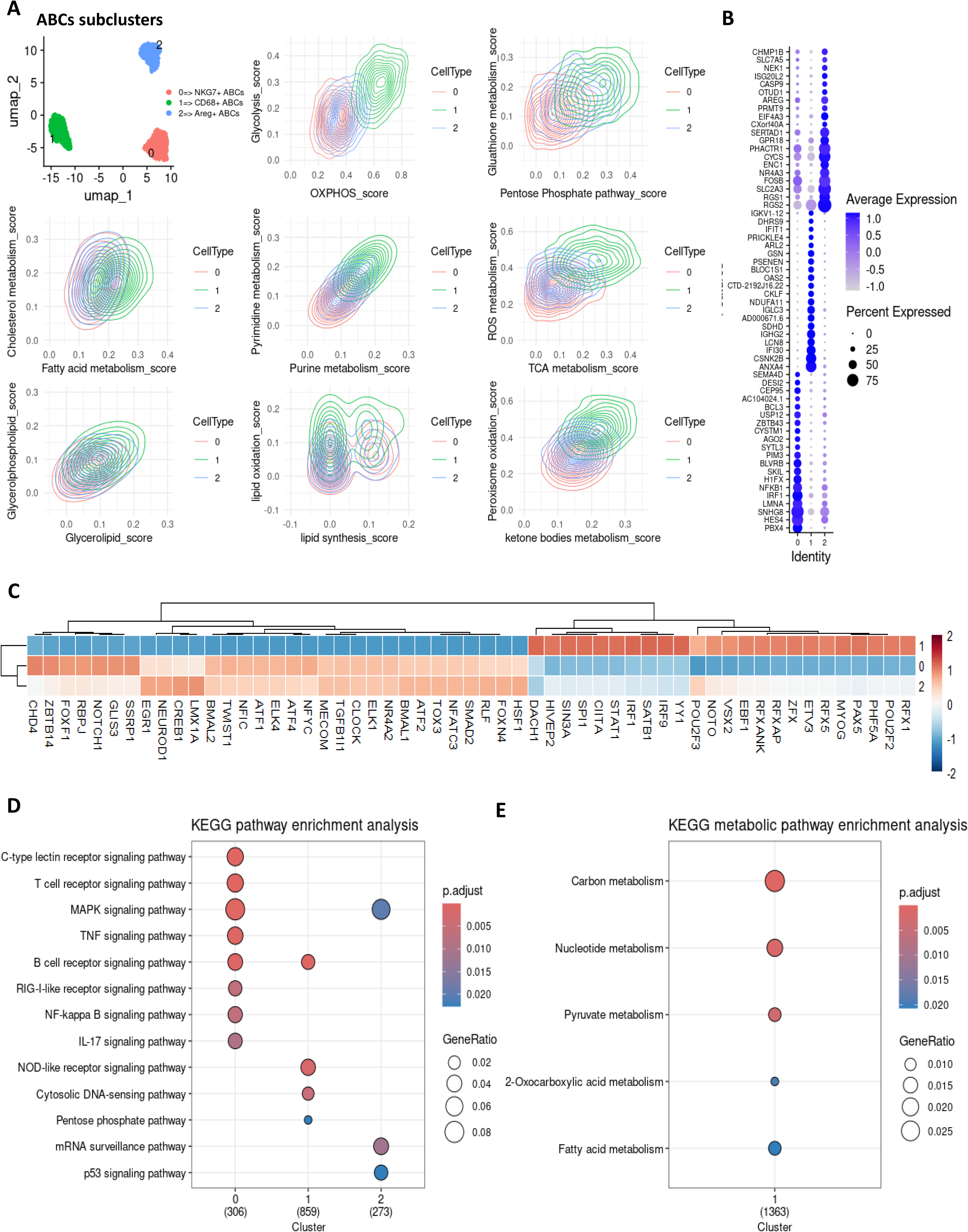
Molecular and metabolic pathway characterization of ABC subclusters (NKG7^+^ ABCs, CD68^+^ ABCs and AREG^+^ ABCs) expanded in the SLE cohort. Pathway and transcriptional analyses were performed to define the functional heterogeneity of ABC subclusters (CellTypes 0=NKG7^+^ ABCs, 1=CD68^+^ ABCs, and 2=AREG^+^ ABCs) that are expanded in individuals with SLE. **(A)** Comparative metabolic pathway analysis using binary plots of selected metabolic programs across ABC subclusters. **(B)** Dot plot showing expression of selected gene markers used to define the immunophenotypic signatures of the three ABC subclusters. **(C)** decoupleR-based transcription factor activity analysis comparing regulatory programs among ABC subclusters. **(D)** Dot plot illustrating KEGG pathway enrichment profiles defining the three ABC subclusters. **(E)** Dot plot showing selectively enriched KEGG metabolic pathways uniquely enriched in ABC subcluster No1.

To gain further insight into the immunological functions of these subpopulations, we performed pathway enrichment analysis using clusterprofiler on the ABC subclusters **(Fig. 4D)**. ABC subcluster 0 was enriched for innate immune and pattern-recognition receptor signalling pathways, including MAPK, TNF, IL-17, and NF-κB signalling, consistent with a strongly pro-inflammatory phenotype. ABC subcluster No1, which exhibited the most hypermetabolic state, showed enrichment of the NOD-like receptor signalling pathway, a key regulator of inflammasome activation and pyroptosis, together with cytosolic DNA-sensing pathways and the NADPH-generating, redox-regulating pentose phosphate pathway. In contrast, ABC subcluster No2, which was shared between patients with SLE and healthy controls, displayed enrichment of p53 signalling and mRNA surveillance pathways, suggestive of a senescent, exhausted, or bystander-like B-cell state **(Fig. 4D)**. Interestingly, only the hypermetabolic ABC subcluster No1, showed unique enrichment of diverse metabolic pathways, such as carbon metabolism, nucleotide metabolism, pyruvate metabolism, 2-oxocarboxylic acid metabolism and fatty acid metabolism **(Fig. 4A, E),** in comparison with the other two ABC subclusters (CellTypes: No0 and No2).

### 3.5 Gene set enrichment and pathway analysis reveals enrichment of phagolysosomal, and immunometabolic programs in lupus ABCs

Gene set enrichment analysis between ABCs and B cells in SLE revealed positive enrichment of the following biological processes: **A)** lysosomal biogenesis, **B)** phagocytosis, **C)** antigen processing and presentation, **D)** systemic lupus erythematosus, **E)** calcium signalling pathways, **F)** metabolic pathways, **G)** sphingolipid metabolism, and **H)** glycerolipid metabolism **(Fig. 5, Supplementary Figure S2).** ABCs in SLE appear to represent a highly activated, metabolically adapted B-cell population with enhanced capacity for antigen uptake, processing, and presentation. Their transcriptional profile suggests they may contribute to lupus pathogenesis by bridging innate-like functions (phagocytosis and lysosomal activity) with adaptive immune activation (antigen presentation and antibody responses).

**Figure 5.**
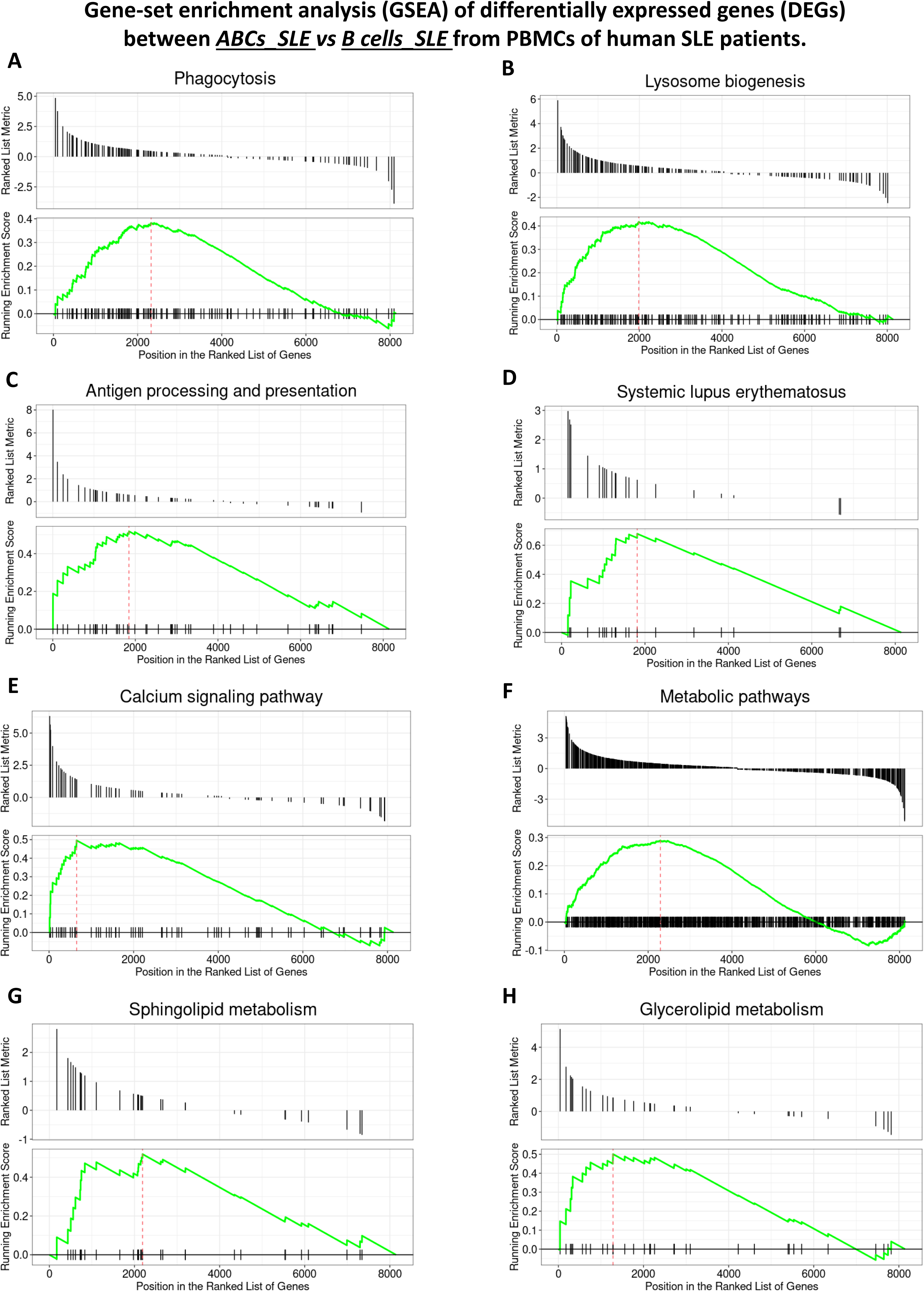
Gene set enrichment analysis (GSEA) of differentially expressed genes (DEGs) between SLE-derived age-associated B cells (ABCs) and SLE-derived B cells. **(A)** Phagocytosis, **(B)** Lysosome biogenesis, **(C)** Antigen processing and presentation, **(D)** Systemic lupus erythematosus, **(E)** Calcium signalling pathway, **(F)** Metabolic pathways, **(G)** Sphingolipid metabolism, **(H)** Glycerolipid metabolism.

A more in-depth KEGG pathway analysis of differentially expressed genes (DEGs) between ABCs and conventional B cells from the SLE patient group, mapped onto the KEGG phagocytosis pathway, revealed gene ontology terms associated with modules relevant to phagosome maturation and phagolysosome formation **(Fig. 6A-B - Supplementary Figure S2)**. More specifically, we observed coordinated upregulation of integrin-mediated and FcγR/Syk/complement (CR3)-dependent adhesion **(Crowley et al., 1997; Jansen et al., 2026)** and uptake machinery, antigen-processing components (MHC class I/II, calnexin, Sec61), and early phagosome maturation factors (Rabex5, Rab5, EEA1, V-ATPase), alongside upregulation of selected degranulation markers (CD63, CD11b, CD13, VAMP2). In contrast, LC3-associated phagocytosis/autophagy-initiation components (Rubicon, VPS34, ATG4, ATG7), the complement receptor CR1, and the proton channel Hv1 were downregulated. A notable shift in NADPH oxidase (NOX2) subunit composition was also observed, with CYBA, NCF2, NCF4, and Rac upregulated but CYBB and NCF1 downregulated across all NOX2 complex instances in the map. Together, these expression changes suggest that ABCs in SLE exhibit enhanced FcγR receptor-mediated phagocytic engagement, antigen-presentation capacity, and early phagolysosomal trafficking, accompanied by altered NADPH oxidase subunit stoichiometry and reduced LC3-associated phagocytosis/autophagic flux relative to conventional B cells **(Fig. 6A)**.

**Figure 6.**
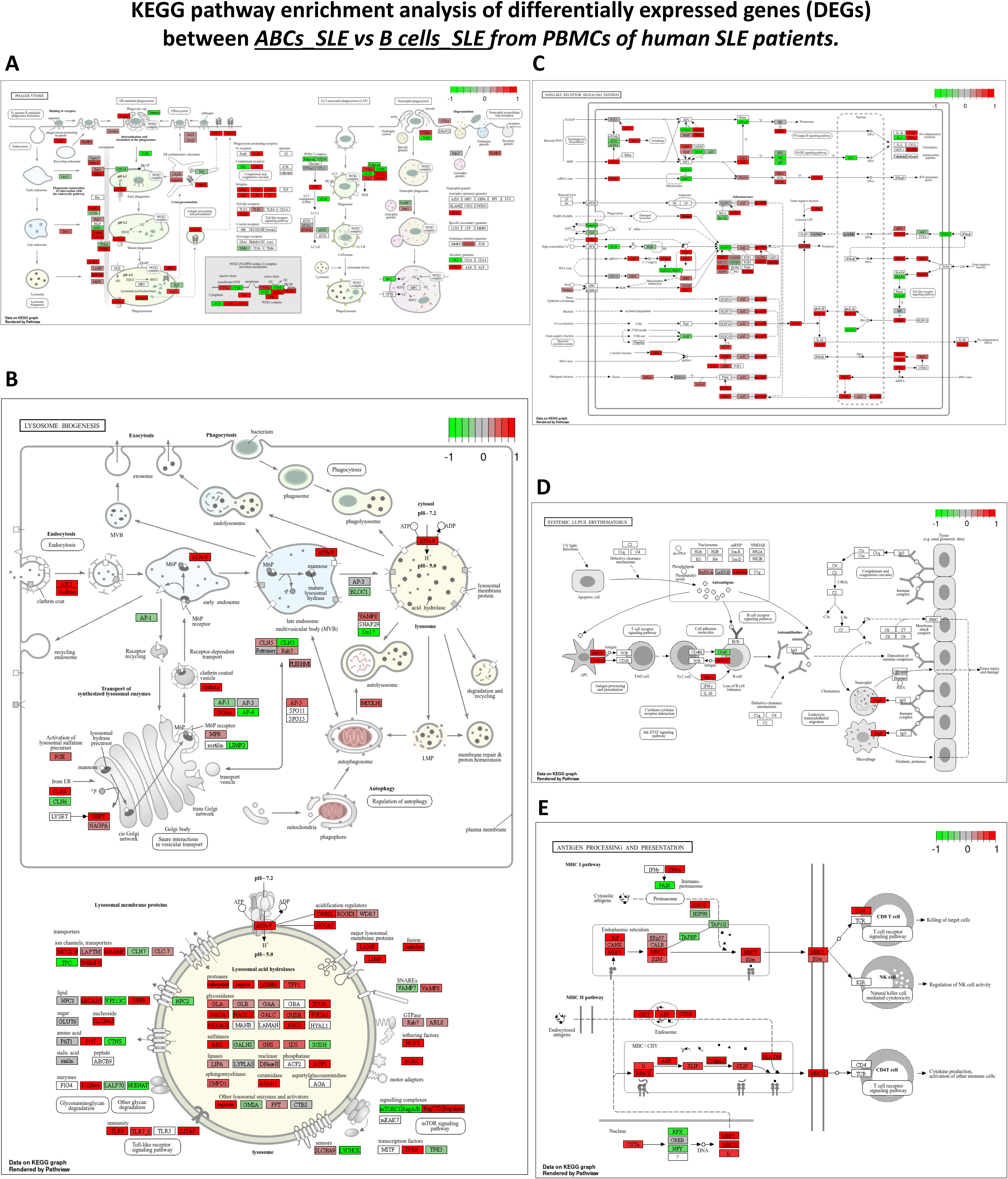
KEGG pathway analysis of DEGs identified between ABCs_SLE and B cells_SLE. Representative enriched KEGG pathways involved in immune regulation and inflammatory responses are shown: **(A)** Phagocytosis, **(B)** Lysosomal biogenesis, **(C)** NOD-like receptor signalling pathway, **(D)** Systemic lupus erythematosus, and **(E)** Antigen processing and presentation. Pathway enrichment indicates distinct immune functional signatures between ABCs_SLE and B cells_SLE.

Further analysis demonstrated substantial activation of the lysosomal pathway in SLE-derived ABCs compared with conventional B cells. Differentially expressed genes were enriched in transcriptional modules governing lysosomal biogenesis, vesicular trafficking, acidification, and proteolytic degradation **(Fig. 6B)**. Notably, multiple vacuolar ATPase subunits, lysosomal membrane proteins, and lysosomal hydrolases were upregulated, indicating enhanced lysosomal activity and degradative capacity. Increased expression of TFEB further suggested activation of the lysosomal biogenesis program. In parallel, elevated TLR7/TLR8-associated signalling components linked lysosomal remodelling with innate immune activation. Collectively, these findings support a metabolically and functionally activated ABC phenotype characterized by augmented intracellular processing and antigen handling **(Fig. 6B)**. Even more importantly, we detected positive enrichment of the NOD-like inflammasome signalling apparatus that activates IL1β and IL18 and initiates pyroptosis through upregulation and assembly of pore-forming gasdermins **(Fig. 6C)**. Upregulation of genes involved in antiviral responses and cytokine production reinforces the consensus that ABCs possess enhanced innate immune sensing capabilities compared with conventional B cells **(Fig. 6C)**.

The systemic lupus erythematosus KEGG pathway showed increased expression of genes involved in antigen presentation and B-cell activation, including CD80/CD86 and MHC class II genes, and molecules associated with T-cell–B-cell interactions **(Fig. 6D)**. Enhanced expression of Fc receptor-related components and immune-complex–associated pathways further supports a role for ABCs in promoting autoreactive immune responses characteristic of lupus **(Fig. 6D)**. Consistent with these observations, and in line with previous studies positioning ABCs in the vicinity of T-B cell borders **(Rubtsov et al., 2015)**, the antigen processing and presentation pathway was strongly enriched in ABCs, compared with the rest of B cells. Multiple genes encoding components of both MHC class I and class II antigen presentation machinery were upregulated, including HLA-DM, CD74, MHC-I, MHC-II, B2M, TAPBP, CALR, CANX, and antigen-processing proteases such as CTSB and AEP. Upregulation of CIITA, a master regulator of MHC class II expression, further indicates enhanced antigen-presenting capacity. Together, these data suggest that ABCs possess a heightened ability to process and present antigens, not only to CD4⁺, but also to CD8⁺ T cells **(Fig. 6E)**.

Overall, the pathway analysis indicates that SLE-associated ABCs are characterized by increased phagolysosomal and inflammatory signalling and enhanced antigen-presentation functions, supporting their proposed role as highly activated immune cells capable of amplifying autoreactive T-cell responses and contributing to lupus pathogenesis. However, further investigation of the functional properties of each ABC subcluster individually suggests that it is the CD68^+^ ABC that is endowed with these properties (e.g., phagocytosis, antigen presentation, lysosomes) **(Supplementary Fig. S3)**.

### 3.6 Interactomics analysis identifies migratory, adhesive and immune-regulatory cell-cell interactions, as well as cholesterol-based metabolite-mediated interaction modes in ABCs

To further substantiate the functional properties of ABCs, we conducted a comprehensive cell-cell interaction analysis using the CellChat package, which models intercellular communication probabilities by incorporating gene expression levels, within an internal curated database of known ligand-receptor pairs and their cofactors **(Fig. 7A & Supplementary Fig. S4)**. Analysis of the interactome showed that ABCs, as sender cells (**Fig. 7B, D)**, exhibited strong inflammatory and antigen-presentation-associated signalling, characterized by prominent MIF–CD74/CD44 and MIF–CD74/CXCR4 interactions as well as extensive HLA class I-mediated communication with CD8^+^ T cells. These findings suggest that ABCs actively participate in shaping both innate and adaptive immune responses. In addition, cell-cell communication analysis displayed predicted ICAM1-dependent interactions consistent with enhanced leukocyte adhesion and potential stabilization of immune synapses **(Fig. 7B, D)**.

**Figure 7.**
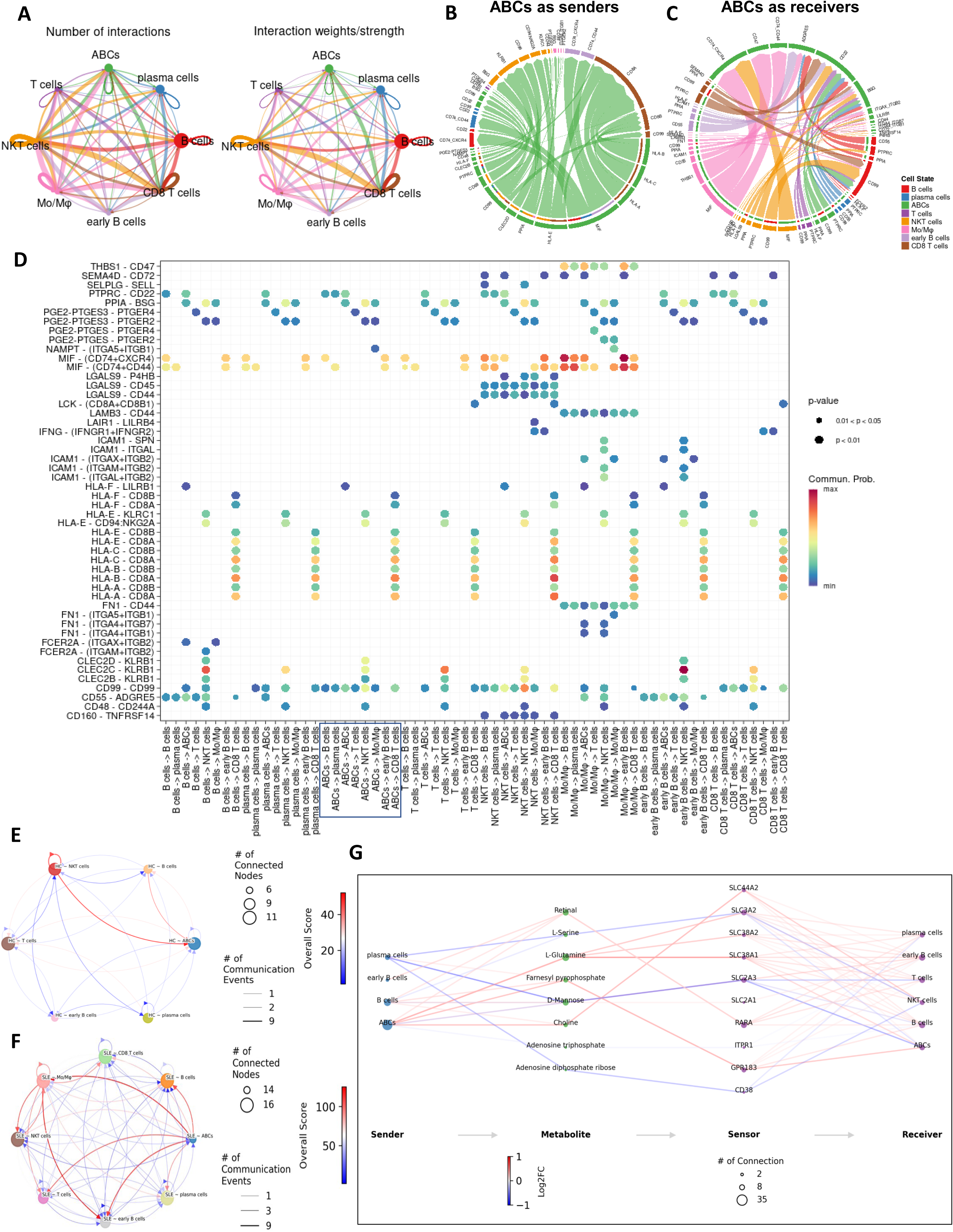
Single-cell transcriptomic and metabolic communication networks reveal altered intercellular interactions in PBMCs from the SLE cohort. Cell–cell communication landscapes were reconstructed using CellChat and MEBOCOST to characterize receptor–ligand and metabolite-mediated signalling networks. **(A)** Schematic workflow of the integrated communication analysis. **(B–C)** Circular plots showing outgoing communication signals from ABCs and incoming signals received by ABCs from other immune cell populations. **(D)** Dot plot summarizing predicted ligand–receptor interaction networks across PBMC cell populations. **(E–F)** Circular plots displaying metabolite-driven communication nodes and their distribution across interacting cell types. **(G)** Differentially altered communication pathways and metabolic interactions associated with ABCs in the SLE cohort.

As receiver cells **(Fig. 7C, D)**, ABCs are targeted by multiple inflammatory and regulatory signalling pathways, including MIF-, prostaglandin E2-, Galectin-9-, and thrombospondin-1-mediated interactions. The coexistence of activating and regulatory signals suggests that ABCs integrate ambivalent activation and functional states. Notably, cell-cell communication analysis also predicted CD22-associated inhibitory signalling in ABCs, suggesting the presence of compensatory or negative feedback regulatory mechanisms that may limit excessive B-cell activation within the inflammatory microenvironment. CD22 is an immunoreceptor with inhibitory motifs, which exerts a suppressive and downmodulating effect on BCR signalling and B-cell activation to maintain immune homeostasis and the resolution of effector responses **(Nitschke, 2009; Waisman et al., 2007)**. Together with PGE2-, LGALS9-, and THBS1-mediated interactions, these findings suggest that ABCs integrate both activating and inhibitory cues, reflecting a wide spectrum of engaging capabilities. Collectively, these results identify ABCs as active participants in the cellular communication network, functioning as both potent signal transmitters and recipients within an inflammatory immune microenvironment **(Fig. 7C, D)**.

It has been shown that, besides cytokines and soluble factors, metabolites constitute important signalling molecules that can facilitate cell-cell communication *(“metabokines”)* **(Tada & Kono, 2025; R. Zheng et al., 2025)**. To examine potential metabolite-driven paracrine communication nodes, we applied MEBOCOST, an established algorithm capable of extrapolating metabolic flux balance analysis from scRNA-seq datasets, so as to detect metabolite-mediated intercellular communication interactions **(R. Zheng et al., 2025)**. Our analysis showed that ABCs were the most active producers of signalling metabolites that potentially modulate and influence other immune cell types, such as T cells and monocytes/macrophages, through various receptors or solute carriers **(Fig. 7E, F)**.

Among the metabolites that are predicted to be biosynthesised by ABCs were retinal, L-Glutamine, farnesyl pyrophosphate and choline. Farnesyl pyrophosphate (FPP) is an important metabolite, which was recently shown to accumulate in migratory dendritic cells (mig-DCs) in SLE-like mice. FPP was found to facilitate not only mig-DC survival but also germinal centre responses in the draining lymph nodes through mitochondrial fusion and increased OXPHOS. In the same study, they applied simvastatin, an HMG-CoA reductase inhibitor of the mevalonate pathway, and the drug reinstated mitochondrial function in mig-DCs and achieved disease remission in SLE mice **(Zhang et al., 2024)**. Our study shows that ABCs could potentially be a source of FPP as well, in line with their hypermetabolic lipid/cholesterol-consuming phenotype **(Fig. 7G)**.

### 3.7 Expansion of Tbet^+^ ABCs in the spleens of female MRL.lpr mice is accompanied by FOXO1 downmodulation

To examine the immunometabolic properties of B cells in an autoimmune mouse model where these ABC cells expand, we used the SLE-like MRL.lpr mouse model. We compared mildly diseased male MRL.lpr to severely diseased female MRL.lpr mice by flow cytometry. Analysis of the splenic B2 B cell subsets revealed tremendous lineage reprogramming from follicular B cells towards marginal zone (MZ) B cells **(Fig. 8A)**. This was accompanied by a significant expansion of Tbet^+^ ABC-like DN B cells **(Fig. 8B),** in line with the literature consensus **(Cancro, 2020; Nickerson et al., 2023; Rubtsova et al., 2015)** .

**Figure 8.**
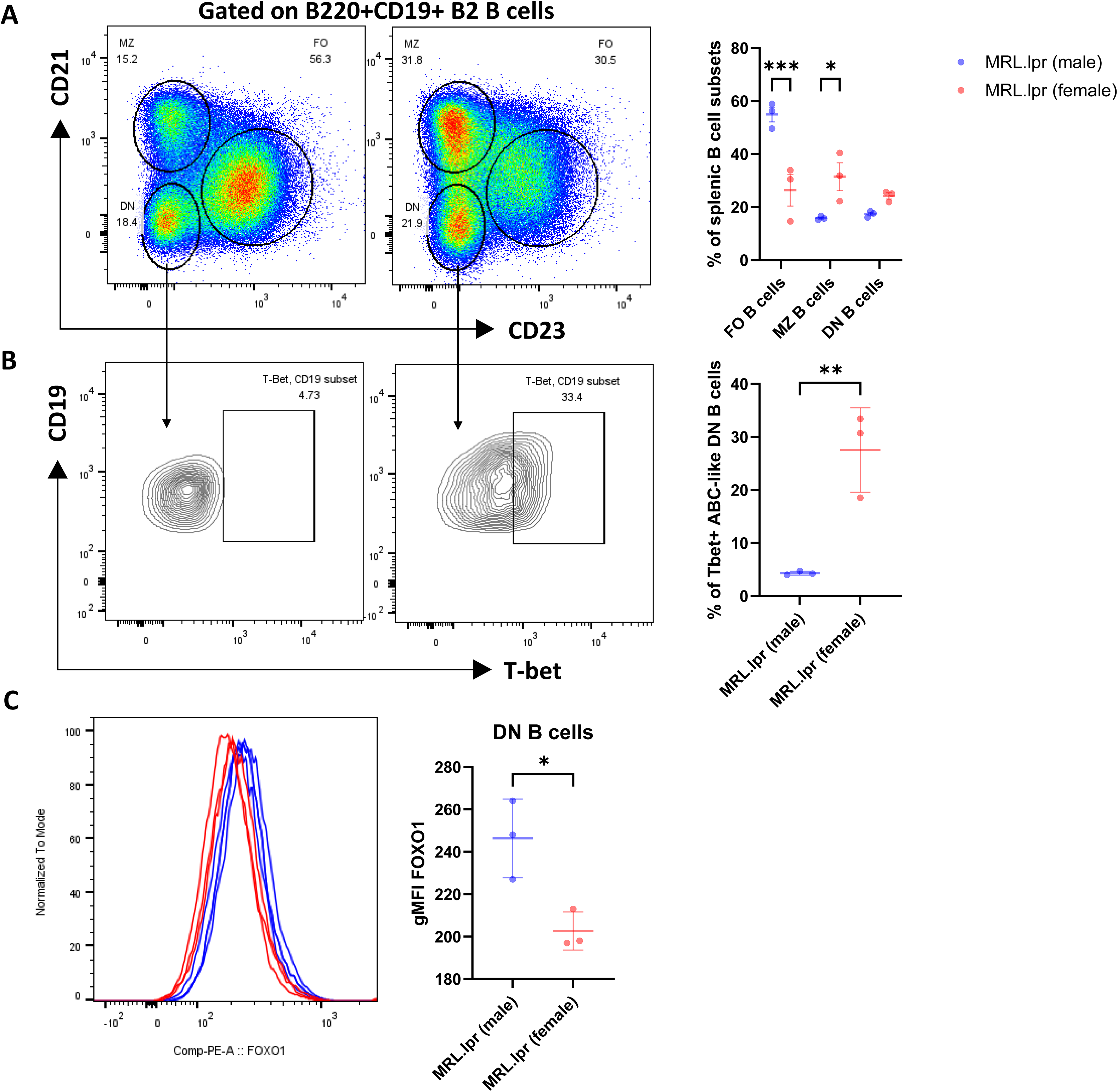
Dissection of B-cell populations in the spleen of male (non-diseased) and female (diseased) MRL.lpr mice. **(A)** Frequencies of follicular (FO), marginal zone (MZ), and double-negative (DN) B-cell populations in the spleen of male and female MRL.lpr mice. **(B)** Frequencies of T-bet⁺ ABC-like DN B cells in male and female MRL.lpr mice. **(C)** Representative histogram showing FOXO1 expression levels in DN B cells from male versus female MRL.lpr mice. Data in graphs are shown as the mean ± SEM and analyzed using two-way ANOVA with Šídák’s multiple comparisons test or two-tailed unpaired Student’s t test. *P < 0.05, **P < 0.01, ***P < 0.001, ****P < 0.0001. Each dot represents one mouse. Data are from at least one independent experiment.

Previously, we demonstrated that constitutive AKT activation biases B cell differentiation towards the marginal zone (MZ) fate through inhibition of FOXO1 **(Cox et al., 2023)**. Interestingly, FOXO1 expression was reduced in the DN B cell population **(Fig. 8C),** consistent with enhanced PI3K-AKT signalling and an activated B cell state. Notably, patients with autoimmune lymphoproliferative syndrome, carrying a loss-of-function mutation in FAS display higher mammalian target of rapamycin (mTOR) **(Staniek et al., 2024)** and AKT signalling compared with healthy controls. Overactive AKT signalling can drive expansion of MZ B cells, through downmodulation of FOXO1 **(Cox et al., 2023)**, which could explain the altered FO/MZ B cell ratio we observe in these mice. Since the MRL.lpr mice similarly carry a loss-of-function mutation in Fas, dysregulated Fas-AKT-mTOR signalling provides a plausible mechanistic framework for the altered FO/MZ B cell ratio observed in these animals.

### 3.8 Comparative scRNA-seq analysis of female MRL.lpr mice validates the hypermetabolic phenotype of ABCs

Previously, **(Nickerson et al., 2023)** reported that ABCs are highly variable with respect to the expression of T-bet, memory markers (PD-L2, CD73, CD80), and plasmablast/antibody-secreting markers (CD138), corresponding to chronically activated cells with the capacity to initiate germinal center reactions and differentiate into plasma cells **(Nickerson et al., 2023).** In the same study **(Nickerson et al., 2023)**, a scRNA/BCR-seq analysis of CD19^+^ B cells from female MRL.lpr mice indicated significant clonal overlap among ABC subsets and plasmablasts, indicative of clonal selection and expansion. We retrieved the CITE-seq dataset GSE202110 from the Gene Expression Omnibus (GEO) repository and performed an immunometabolic analysis to characterize metabolic differences among CD19^+^ B cell populations and across their differentiation states.

Murine ABCs from the MRL.lpr strain were identified by the lack of Cr2 (CD21) and Fcer2a (CD23) expression **(Fig. 9A, B)**. To ensure consistency with the CITE-seq dataset, we benchmarked the ABC populations detected via flow cytometry, which faithfully captured the phenotypic heterogeneity within the ABC compartment as illustrated in the CITE-seq dataset **(Fig. 9C, D)**.

**Figure 9.**
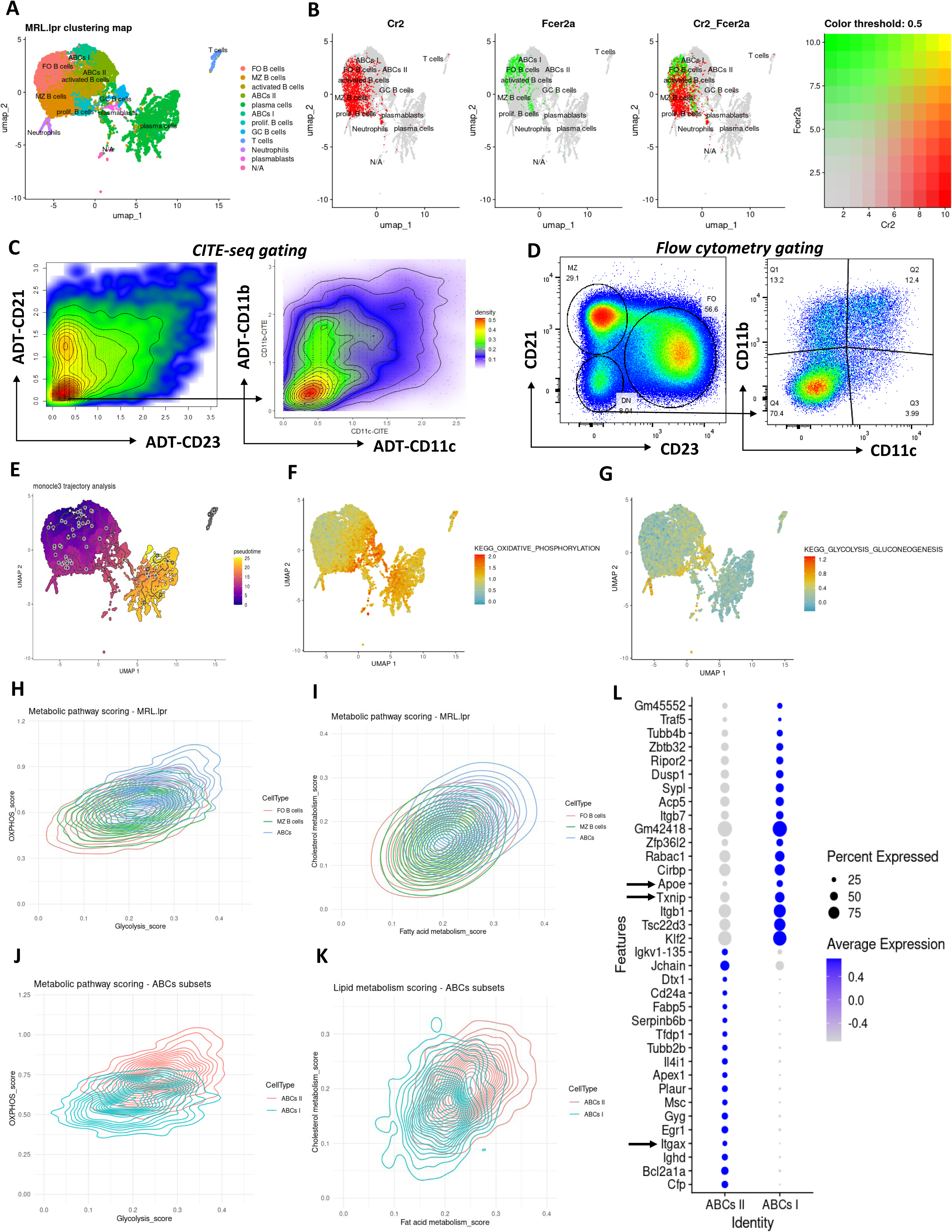
Single-cell transcriptomic re-analysis of splenic CD19⁺ B cells from female MRL.lpr mice reveals distinct ABC-like DN B-cell states. Single-cell RNA sequencing (scRNA-seq) re-analysis was performed on splenic CD19⁺ B cells isolated from three female MRL.lpr mice to characterize B-cell heterogeneity and metabolic states, from the GSE202110 dataset. **(A)** UMAP visualization of the splenic CD19⁺ B-cell population. **(B)** Identification and characterization of CD21⁻CD23⁻ double-negative (DN) B cells. **(C–D)** CITE-seq-based annotation of B-cell clusters, including follicular (FO) B cells, marginal zone (MZ) B cells, and CD21⁻CD23⁻ ABC-like DN B cells, with validation by flow cytometry. **(E)** Monocle3 trajectory analysis revealing developmental relationships among B-cell subsets. **(F–G)** UMAP visualization of pathway activity scores for KEGG oxidative phosphorylation and KEGG glycolysis/gluconeogenesis pathways. **(H–I)** Metabolic pathway scoring of FO B cells, MZ B cells, and ABCs based on oxidative phosphorylation (OXPHOS) versus glycolysis scores and cholesterol metabolism versus fatty acid metabolism scores. **(J–K)** Comparative metabolic pathway scoring of ABC subclusters (ABCs I and ABCs II) based on OXPHOS versus glycolysis and cholesterol versus fatty acid metabolism signatures. **(L)** Phenotypic fingerprinting of the two ABC subclusters (ABCs I and ABCs II).

Again, we conducted a pseudotime trajectory analysis with Monocle3 to map differentiation trajectory lines, which showed a clear trajectory from naïve/immature B cells to ABCs, and from there to plasma cells (**Fig. 9E)**. Along this line, we observed an increase in OXPHOS and glycolysis among the ABCs (**Fig. 9F, G)**. Pathway scoring showed that the ABCs represented the metabolically most active population among the B2 B cell family, in terms of OXPHOS, glycolysis, and lipid utilization **(Fig. 9H, I)**. Within the ABC group, we could clearly detect 2 subsets (annotated as “ABCs I” and “ABCs II”), which exist in two metabolic states, in terms of the same metabolic statuses **(Fig. 9J, K)**. The metabolically active subset (ABCs II) corresponds to CD11c^+^ ABCs, while the other one (ABCs I) belongs to an apolipoprotein E (ΑpoE^+^)-positive subset, wherein apolipoprotein E potentially fuels lipid metabolism **(Fig. 9L)**.

### 3.9 SCENITH reveals metabolic reprogramming and reduced protein synthesis in CD21^-^CD23^-^ B cells from severely diseased female MRL.lpr mice

To substantiate the scRNA-seq results, we employed SCENITH, a flow cytometry-based metabolic assay that tries to infer metabolic dependencies by measuring protein translation levels, through puromycin incorporation into nascently synthesised protein chains by the ribosomes, across different treatment states. We compared protein puromycinylation levels between mildly diseased male and severely diseased female MRL.lpr mice in different splenic B cell subsets. In the male group, inhibition of OXPHOS with oligomycin (ATPase V inhibitor) revealed a bimodal puromycin distribution **(Fig. 10A, blue and red)**. Backgating of the puromycin peaks onto the CD21 versus CD23 distribution, revealed profound metabolic heterogeneity **(Fig 10A-D).** In mildly-diseased male MRL.lpr mice, oligomycin-treatment of spleen cells ex-vivo revealed two puromycin peaks **(Fig. 10A-B).** The puromycin-low peak, which is reflective of an OXPHOS-dependent state, corresponded mostly to CD21^-^CD23^-^ ABC-like DN B cells, while the puromycin-high peak, which is reflective of a glycolytic state, belonged mostly to marginal zone B cells and follicular B cells **(Fig. 10A-B)**.

**Figure 10.**
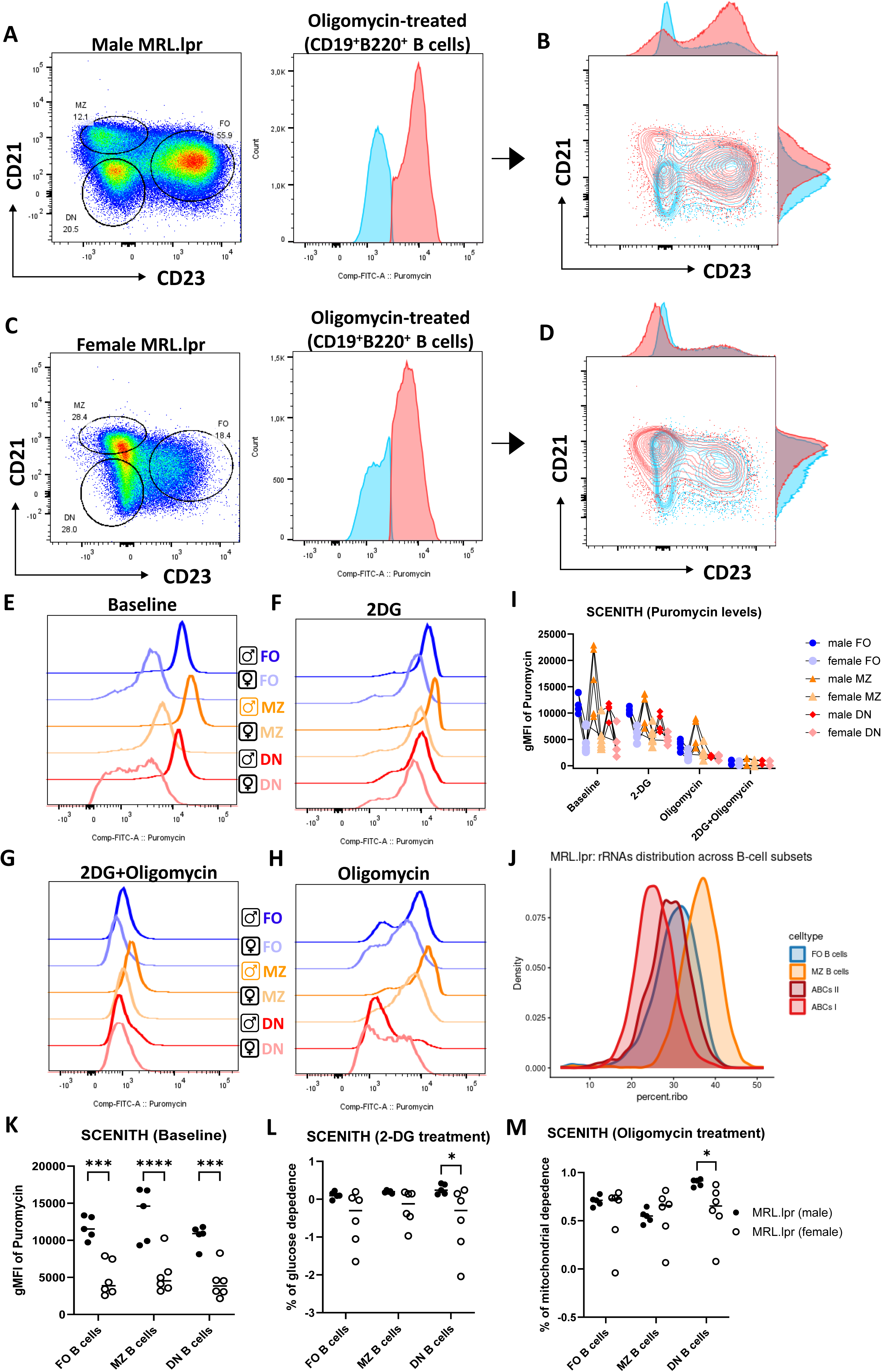
SCENITH analysis reveals distinct metabolic dependencies of B-cell subsets in male (non-diseased) and female (diseased) MRL.lpr SLE-like mice. The cellular metabolic state of splenic B cells from male and female MRL.lpr mice was assessed using the SCENITH assay to quantify translation activity, glucose dependence, and mitochondrial dependence. **(A–B)** Representative puromycin incorporation histograms showing the bimodal distribution of total B2 B cells following oligomycin treatment of splenocytes from male MRL.lpr mice, with corresponding backgating analysis on CD21 versus CD23 expression to identify B-cell subsets. **(C–D)** Representative puromycin incorporation histograms showing the bimodal distribution of total B2 B cells following oligomycin treatment of splenocytes from female MRL.lpr mice, with corresponding CD21 versus CD23 backgating analysis. **(E–H)** Overlay of baseline puromycin incorporation profiles from follicular (FO), marginal zone (MZ), and double-negative (DN) B cells from male and female MRL.lpr mice under different metabolic perturbations: (E) baseline conditions, (F) 2-deoxy-D-glucose (2DG) treatment, (G) combined 2DG and oligomycin treatment, and (H) oligomycin-only treatment. **(I)** Quantification of SCENITH-derived metabolic parameters across B-cell subsets. **(J)** Cross-validation of SCENITH-based metabolic activity measurements with scRNA-seq-derived ribosomal RNA (rRNA) expression levels at steady state in female MRL.lpr mice. **(K–M)** Quantification of (K) puromycin incorporation levels at steady state, (L) glucose dependence, and (M) mitochondrial dependence across B-cell populations. Data in graphs are shown as the mean ± SEM and analyzed using two-way ANOVA with Šídák’s multiple comparisons test or two-tailed unpaired Student’s t test. *P < 0.05, **P < 0.01, ***P < 0.001, ****P < 0.0001. Each dot represents one mouse. Data are pooled together from two independent experiments (n=5-6).

In the severely-diseased female MRL.lpr group, a proportion of marginal zone B cells exhibited increased OXPHOS activity, together with the CD21⁻CD23⁻ ABC-like B-cell population **(Fig. 10C–D)**. In contrast, baseline puromycinylation levels were significantly lower across all B-cell subsets in female mice compared with their corresponding populations in the male cohort **(Fig. 10E, I, K)**, indicating an overall reduction in basal protein synthesis activity in female-derived B cells. Notably, the CD21⁻CD23⁻ DN B-cell population, in the female cohort, displayed a bimodal distribution of puromycin incorporation at baseline, with a subset of cells exhibiting relatively high translational activity and another subset showing markedly lower activity. This bimodal pattern was consistent with differences in ribosomal gene expression observed in the MRL.lpr scRNA-seq dataset **(Fig. 10E, I, J)**. These findings suggest that metabolic and translational activity may provide additional parameters for resolving functionally distinct subpopulations within the phenotypically heterogeneous ABC compartment.

Further evidence for metabolic heterogeneity among ABCs was obtained following oligomycin treatment. In the female cohort, ABCs also displayed a bimodal distribution of puromycin incorporation, indicating that these cells differ in their ability to sustain protein synthesis when OXPHOS is blocked. This pattern is consistent with the presence of at least two metabolically distinct fractions: an OXPHOS-dependent population, whose translational activity is reduced upon mitochondrial inhibition, and a second population that is better able to maintain activity through glycolytic or other compensatory metabolic pathways. The latter profile is in agreement with the metabolic heterogeneity inferred from the scRNA-seq datasets generated from the MRL.lpr strain **(Fig. 10H)**. Together, these observations support the notion that the ABC compartment is not metabolically uniform, but instead comprises subpopulations with distinct bioenergetic dependencies.

In summary, our SCENITH analyses reveal metabolic versatility, with enhanced mobilization of both glycolytic and mitochondrial respiratory pathways in CD21⁻CD23⁻ DN B cells from the female, severely diseased MRL.lpr mice. Following 2-DG treatment, CD21⁻CD23⁻ B cells from female MRL.lpr mice exhibited a relative shift toward increased respiratory capacity compared with their male counterparts, suggesting that these cells may have a greater capacity to maintain energy production through substrates other than glucose, such as amino acids or lipids. In addition, the persistence of puromycin incorporation following oligomycin treatment indicates that a fraction of these cells can maintain translational activity despite inhibition of mitochondrial ATP production, consistent with an increased capacity to engage extra-mitochondrial sources of energy. Thus, rather than relying predominantly on a single metabolic pathway, CD21⁻CD23⁻ DN B cells from female MRL.lpr mice display metabolic flexibility and adaptability. This metabolic flexibility may contribute to the ability of these cells to sustain cellular functions under conditions of altered nutrient availability and mitochondrial inhibition and provides an additional layer of heterogeneity that complements the transcriptional differences identified by scRNA-seq **(Fig. 10F, H, L, M)**.

### 3.10 ABCs from MRL.lpr are enriched for ubiquitination, phagocytosis, and antigen-processing & presentation ontology terms

Gene ontology analysis with the clusterprofiler package, showed representation of terms associated with protein ubiquitination, particularly in the hypermetabolic (ABCs II) subset, which aligns with the SCENITH-depicted phenotype of lower protein levels in the female MRL.lpr cohort **(Fig. 11A)**. In fact, both FO and MZ B cells from the female MRL.lpr mice were enriched for ontology terms associated with ubiquitination **(Fig. 11A).** KEGG pathway analysis for ABCs showed upregulation of various pathways including endocytosis, PD-L1 expression, protein processing in the endoplasmic reticulum, antigen processing & presentation, phagosome formation and endocytosis, ROS production, as well as profound disease-term enrichment in pathways associated with inflammation and proteotoxicity **(Fig. 11B-C)**.

**Figure 11.**
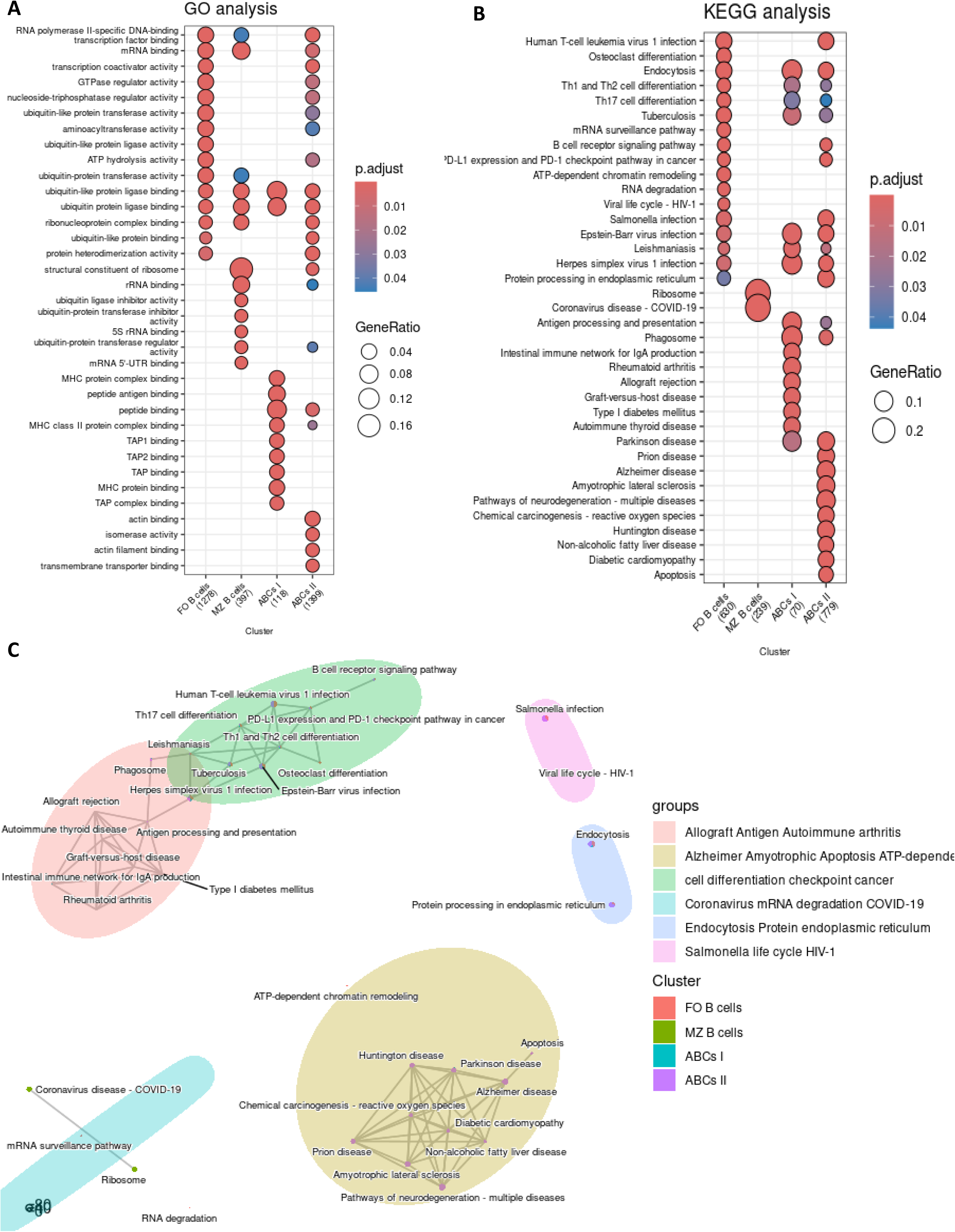
Transcriptomic pathway signatures distinguish ABCs from conventional FO and MZ B cells in female MRL.lpr mice. Differentially expressed genes between ABCs and FO/MZ B cells were analyzed using gene set enrichment analysis (GSEA) and KEGG pathway enrichment. **(A–B)** GSEA plots depicting enrichment of TGF-β signalling and p53 signalling pathways in ABCs. **(C)** KEGG pathway visualization of genes upregulated in ABCs compared with FO and MZ B cells.

### 3.11 Pathway analysis reveals distinct functional programs in ABCs relative to FO and MZ B cells among female MRL.lpr mice

Gene set enrichment analysis revealed relative overrepresentation of the TGF-b and p53 signalling pathways among the ABCs, relative to FO/MZ B cells **(Fig. 12A, B)**. The p53 pathway is associated with a state of immunosenescence **(Fu et al., 2025)**, while the TGF-β factor is important for immune homeostasis, and is commonly classified as an immunosuppressive cytokine as it functions to suppress the activity of pro-inflammatory cells while promoting the expansion of immunosuppressive cells **(Deng et al., 2024)**.

**Figure 12.**
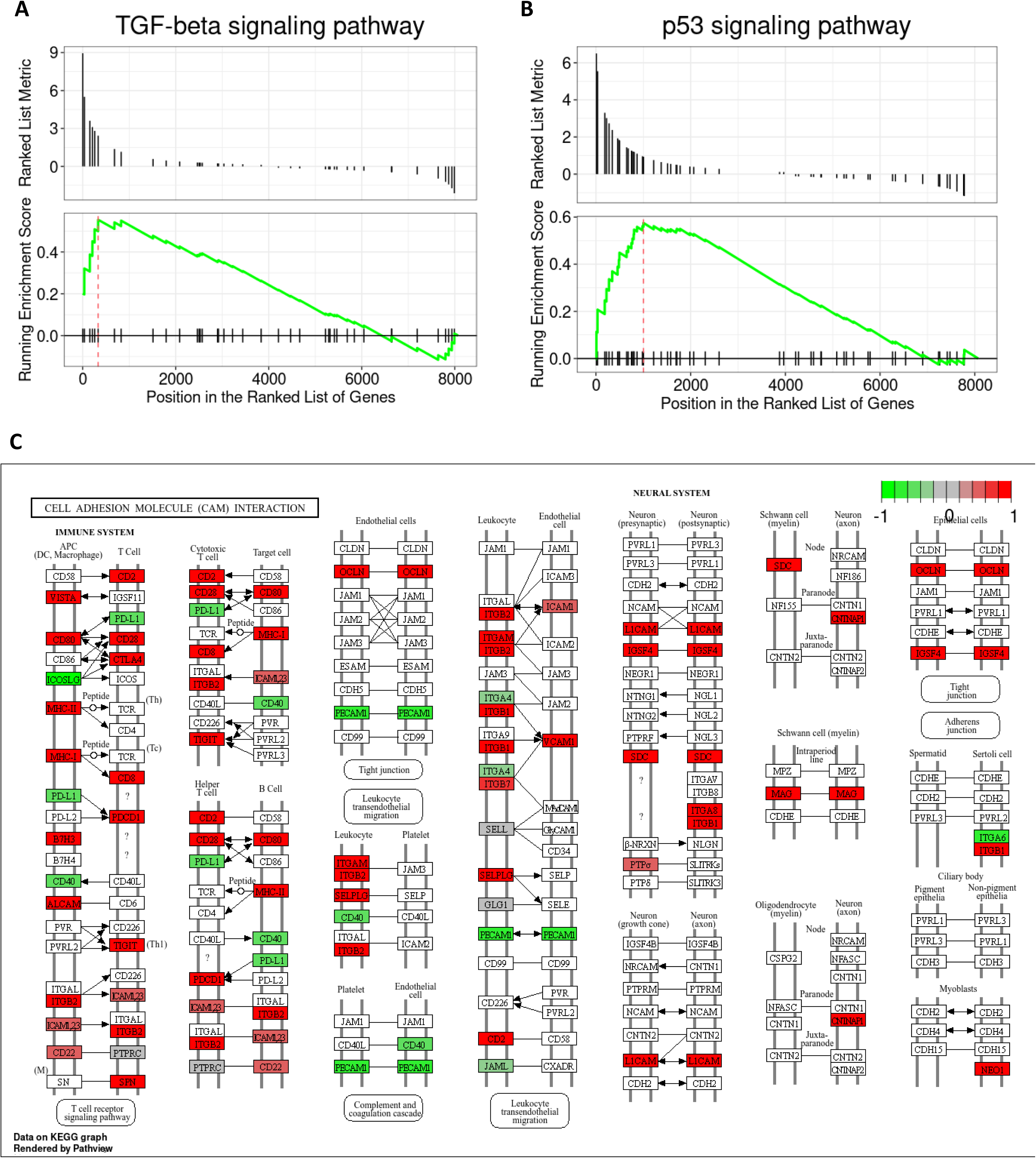
ClusterProfiler-based functional enrichment analysis of B2 B-cell clusters from the scRNA-seq dataset of female MRL.lpr mice. Functional annotation and pathway analysis were performed on B2 B-cell clusters identified from the splenic scRNA-seq dataset of female MRL.lpr mice using ClusterProfiler. **(A)** Dot plot showing gene ontology (GO) enrichment terms across distinct B2 B-cell populations. **(B)** Dot plot displaying enriched Kyoto Encyclopedia of Genes and Genomes (KEGG) pathways across B2 B-cell populations. **(C)** Network visualization of enriched GO terms grouped according to semantic similarity, highlighting functional relationships among biological processes.

The “KEGG-Cell Adhesion Molecule (CAM)” interaction pathway analysis revealed extensive transcriptional remodelling in age-associated B cells (ABCs) compared with follicular/marginal zone (FO/MZ) B cells from female MRL.lpr mice **(Fig. 12C)**. Differentially expressed genes (DEGs) were predominantly enriched in immune cell adhesion and co-stimulatory networks, with marked upregulation of molecules involved in T cell–B cell interactions, antigen presentation, and leukocyte trafficking **(Fig. 12C)**. ABCs exhibited increased expression of key immune checkpoints, and regulatory-like or exhaustion markers, including CTLA4, PDCD1 (PD-1), CD2, TIGIT, ICAM1, ICAM2, ITGB1, ITGB2, SELPLG (Selectin-P ligand), and MHC-I/MHC-II–associated molecules, suggesting enhanced capacity for immune synapse formation and cellular communication. In contrast, several adhesion molecules such as PECAM1, PD-L1, ITGA4, and ITGA6, or costimulatory molecules like CD40, were relatively downregulated. Beyond immune-specific interactions, altered expression of cell adhesion molecules involved in endothelial adhesion, trans-endothelial migration, and epithelial/neural adhesion modules indicates broader changes in cell–cell interaction programs. Collectively, and in line with our prior patient scRNA-seq analysis, these findings demonstrate that ABCs, in the MRL.lpr context, acquire a mixed phenotype by combining stimulatory signalling, integrin-mediated adhesion, and leukocyte trafficking pathways, but also regulatory-like or immunomodulatory features **(Fig. 12C)**.

### 3.12 Transcriptomics-based in silico drug-repurposing analysis highlights HMG-CοA reductase inhibitors and proteasome inhibitors as potential options for SLE management

For translational purposes, we implemented a variety of standard computational tools and pipelines (DMEA, SCANet, clusterprofiler, and ASGARD) to conduct in silico drug repurposing analysis to inquire how pathway perturbations could potentially be exploited for therapeutic purposes, by repositioning already approved drug options. We generated lists of differentially expressed genes for ABCs and B cells, between SLE patients and healthy individuals (HC), which were used for drug mechanism enrichment analysis using the DMEA tool. Our analysis suggested several druggable targets that could exert an impact on ABCs, including RAF inhibitors, AP inhibitors, HMGCR inhibitors, MEK inhibitors, and antioxidants **(Fig. 13A)**.

**Figure 13.**
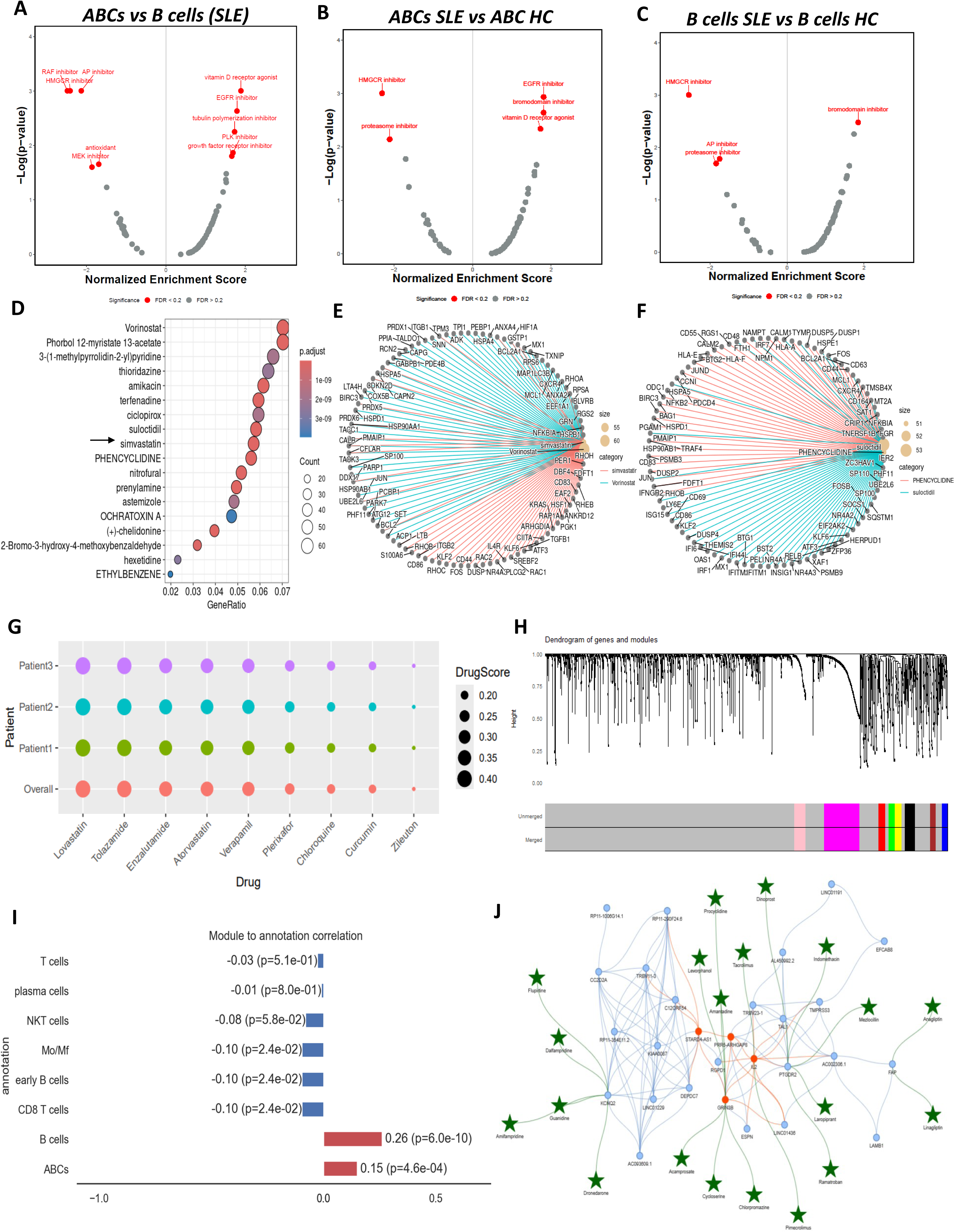
Computational drug repurposing identifies candidate compounds targeting SLE-associated B-cell transcriptional programs. Transcriptomic signatures from SLE B-cell populations were integrated with drug perturbation databases and computational pipelines to identify potential therapeutic interventions. **(A–C)** Drug mechanism enrichment analysis using DEGs from ABCs versus B cells in SLE, ABCs from SLE versus healthy controls, and B cells from SLE versus healthy controls. **(D)** ClusterProfiler-based drug enrichment analysis showing significantly associated compounds. **(E–F)** Drug–gene interaction networks highlighting predicted targets of selected compounds, including simvastatin, vorinostat, phencyclidine, and suloctidil. **(G–H)** Drug repurposing predictions generated using ASGARD and SCAnet pipelines. **(I–J)** Identification of candidate drugs predicted to modulate transcriptional modules enriched in B cells and ABCs.

Both ABCs and B cells in general, from the SLE patients, appear to be vulnerable to HMGCR and proteasome inhibitors **(Fig. 13B-C)**. In light of the hypermetabolic OXPHOS-driven, potentially lipid-fuelled phenotype of ABCs, coupled to the lower protein output, detected by SCENITH, we postulate that hyperactive B cells face high risk of proteotoxic risk, perhaps due to the accumulation of misfolded proteins. Proteasome inhibitors have been used to treat plasmacytosis in autoimmune diseases, and also in the cancer context, such as in multiple myeloma **(Gomez et al., 2011; Neubert et al., 2008)**. Drug-set enrichment analysis **(Fig. 13D-F)** and the ASGARD pipeline **(Fig. 13G)** also recommended statins as a useful approach to target cholesterol metabolism, since cholesterol and other lipid metabolites were found to be mobilized by ABCs.

Finally, drug-repurposing with SCANet identified a transcriptional module significantly associated with ABCs **(Fig. 13H-I).** Candidate drugs included compounds such as tacrolimus, indomethacin, pimecrolimus, chlorpromazine, and amantadine, among others. Based on the ABC-associated module identified by SCANet, we prioritized candidate drugs according to their known immunomodulatory mechanisms. Tacrolimus and pimecrolimus emerged as particularly compelling candidates because both inhibit FKBP12–calcineurin signalling, a pathway capable of regulating inflammatory cytokine transcription and lymphocyte activation **(Nghiem et al., 2002)**. Indomethacin, ramatroban, and laropiprant were prioritized as complementary candidates targeting prostaglandin-/thromboxane-mediated inflammatory signalling. Together, these candidates represent two mechanistically distinct therapeutic axes—calcineurin-dependent immune signalling and eicosanoid signalling—that may modulate the ABC-associated transcriptional program **(Fig. 13J)**.

### 3.13 ABCs and inflammaging: Mitochondrial parameter evaluation shows that ABCs are undergoing mitochondrial remodeling in aging

Next, we assessed mitochondrial mass and membrane potential using the MitoTracker Green and TMRE dyes. CD21^-^CD23^-^ DN B cells exhibited elevated mitochondrial membrane potential, but not increased mitochondrial mass **(Fig. 14A)**. Further normalization of TMRE intensity to mitochondrial content (gMFI_TMRE/_gMFI_MitoTracker r_atio) indicated increased membrane potential per mitochondrion **(Fig. 14A)**. However, plotting total CD19^+^B220^+^ B2 B cells on a MitoTrackerGreen vs SSC-A binary plot, revealed two subsets, with expansion of hypergranular and hypermitochondrial SSC-A^HIGH^Mitotracker^HIGH^ B cells in the aged cohort compared to young mice (**Fig. 14B).** SSC-A^HIGH^Mitotracker^HIGH^ B cells were enriched for CD21^-^CD23^-^ DN B cells, at ranges of 10-20%, suggesting metabolic heterogeneity within the CD21^-^CD23^-^ DN B cell compartment. Furthermore, the hypergranular (SSC-A^high^) fraction of CD21^-^CD23^-^ DN B cells displayed high levels of NADH and increased accumulation of the voltage-driven TMRE dye, altogether indicating altered cellular redox/metabolic state and mitochondrial polarization **(Fig. 14B-D)**. In summary, in the aged cohort, when evaluating Mitochondrial mass (MitotrackerGreen) in correlation with SSC-A, we clearly observe 2 subpopulations, one SSCA^low^MitoTracker^low^NADH^+^TMRE^+^ and one SSCA^high^MitoTracker^high^NADH^high^TMRE^high^ subset **(Fig. 14C-D).** Thus, the linkage between metabolic and mitochondrial activity, and granularity constitutes a demarcating layer that could be used to distinguish phenotypic heterogeneity within the CD21^-^CD23^-^ DN B cell fraction.

**Figure 14.**
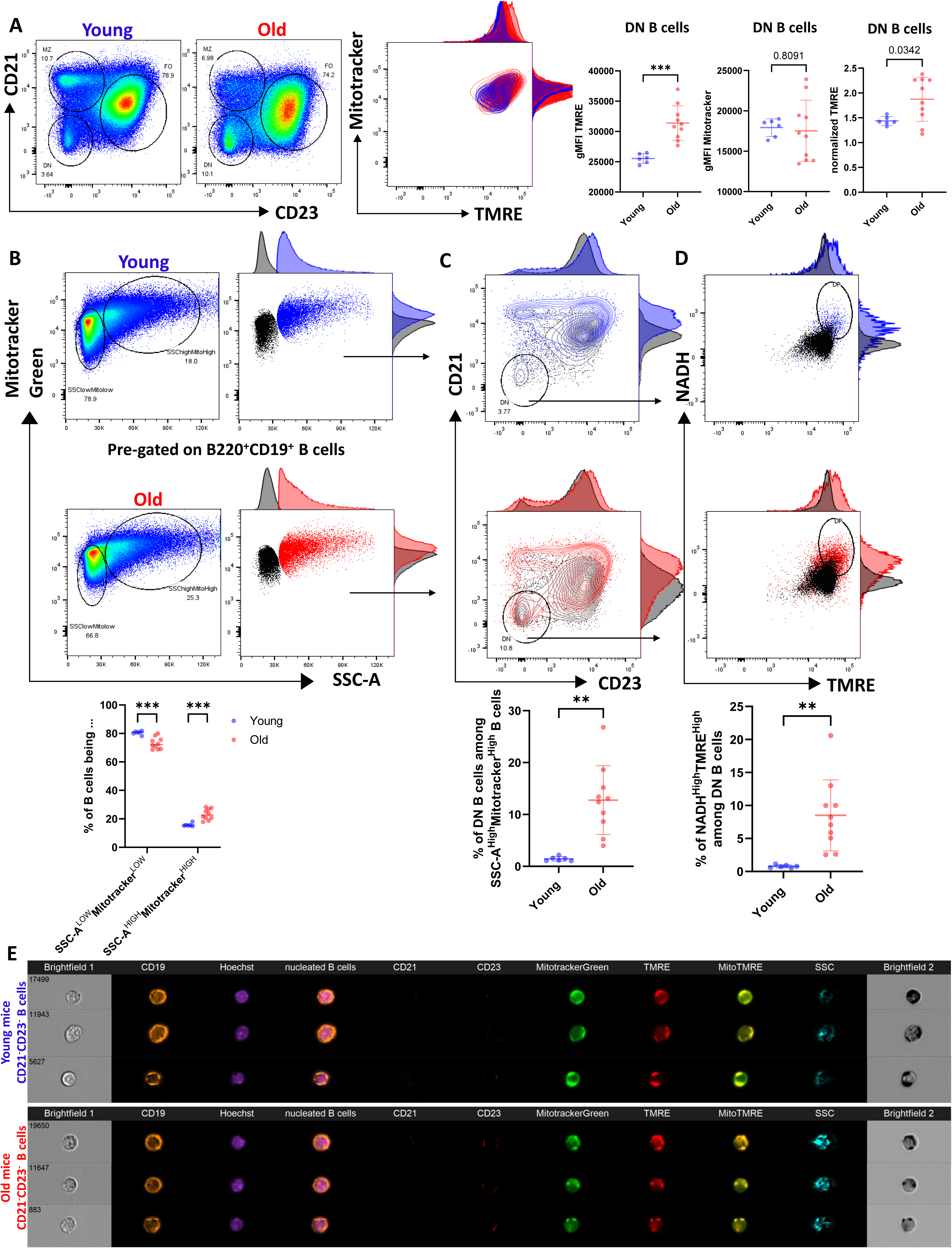
Assessment of mitochondrial mass and membrane potential in B-cell subsets using MitoTracker Green and TMRE staining. Mitochondrial content and function were evaluated in splenic B cells from young and aged mice using MitoTracker Green and tetramethylrhodamine ethyl ester (TMRE) staining to assess mitochondrial mass and membrane potential, respectively. **(A)** Evaluation of MitoTracker Green and TMRE levels in CD21⁻CD23⁻ DN B cells in young and aged mice. **(B)** Identification of hypergranular, hypermitochondrial SSCA^high^MitoTracker^high^ CD19⁺B220⁺ B2 B cells by flow cytometry. **(C)** Backgating of SSCA^high^MitoTracker^high^ B2 B cells onto CD21 versus CD23 expression plots to determine their distribution among follicular (FO), marginal zone (MZ), and double-negative (DN) B-cell subsets. **(D)** Backgating of DN B cells on NADH versus TMRE plots to assess metabolic state and mitochondrial membrane potential. **(E)** Spatial visualization with imaging flow cytometry of MitoTracker Green, TMRE and side-scatter granularity in CD21-CD23-ABC-like B cells from young vs old mice. Data in graphs are shown as the mean ± SEM and analysed using two-tailed unpaired Student’s t test. *P < 0.05, **P < 0.01, ***P < 0.001, ****P < 0.0001. Each dot represents one mouse (n=6-10).

Mitochondrial visualization with imaging flow cytometry confirmed increased granularity spots (SSC-A), among the CD21^-^CD23^-^ B cells in aged versus young mice, which could partially be attributed to mitochondrial mass **(Fig. 14E)**. Together, these data indicate that the heightened metabolic state of aged ABCs is primarily driven by increased mitochondrial remodelling or perhaps increased consumption of energy-dense lipids, whose oxidation generated a vast number of electrons pumped into the mitochondrial matrix. This mitochondrial adaptation may provide the energetic and biosynthetic foundation required to sustain the inflammatory and autoreactive phenotype characteristic of ABCs.

### 3.14 ABCs and inflammaging: Redox balance recalibration of ABCs in young and old mice reveals a hypermetabolic phenotype specific to ABCs

To delineate how redox metabolic dynamics are altered during aging, with particular emphasis on the CD21⁻CD23⁻CD19⁺B220⁺ DN B-cell compartment, we performed comprehensive NADH/FAD redox profiling in splenic B cells isolated from young and aged mice. Conventional measurements such as oxygen consumption rate (OCR) represent an aggregate readout of cellular oxygen utilization and do not provide insight into mitochondrial abundance, individual mitochondrial activity, or the underlying redox state driving metabolic flux **(Abir et al., 2024; Lemire et al., 2022; Stylianakis & Hövelmeyer, 2026)**.

Following exclusion of T cells, myeloid cells, and B1 B cells, we quantified real-time NADH and FAD fluorescence kinetics and derived the optical redox ratio during a Seahorse-inspired mitochondrial stress assay on B2 B cell subsets **(Fig. 15, Supplementary Figure S6)**. Among the analysed B-cell populations, as illustrated in the gating strategy **(Supplementary Figure S6 A)**, double-negative (DN) B cells exhibited the most pronounced metabolic remodelling **(Supplementary Figure S6 B-E)**.

**Figure 15.**
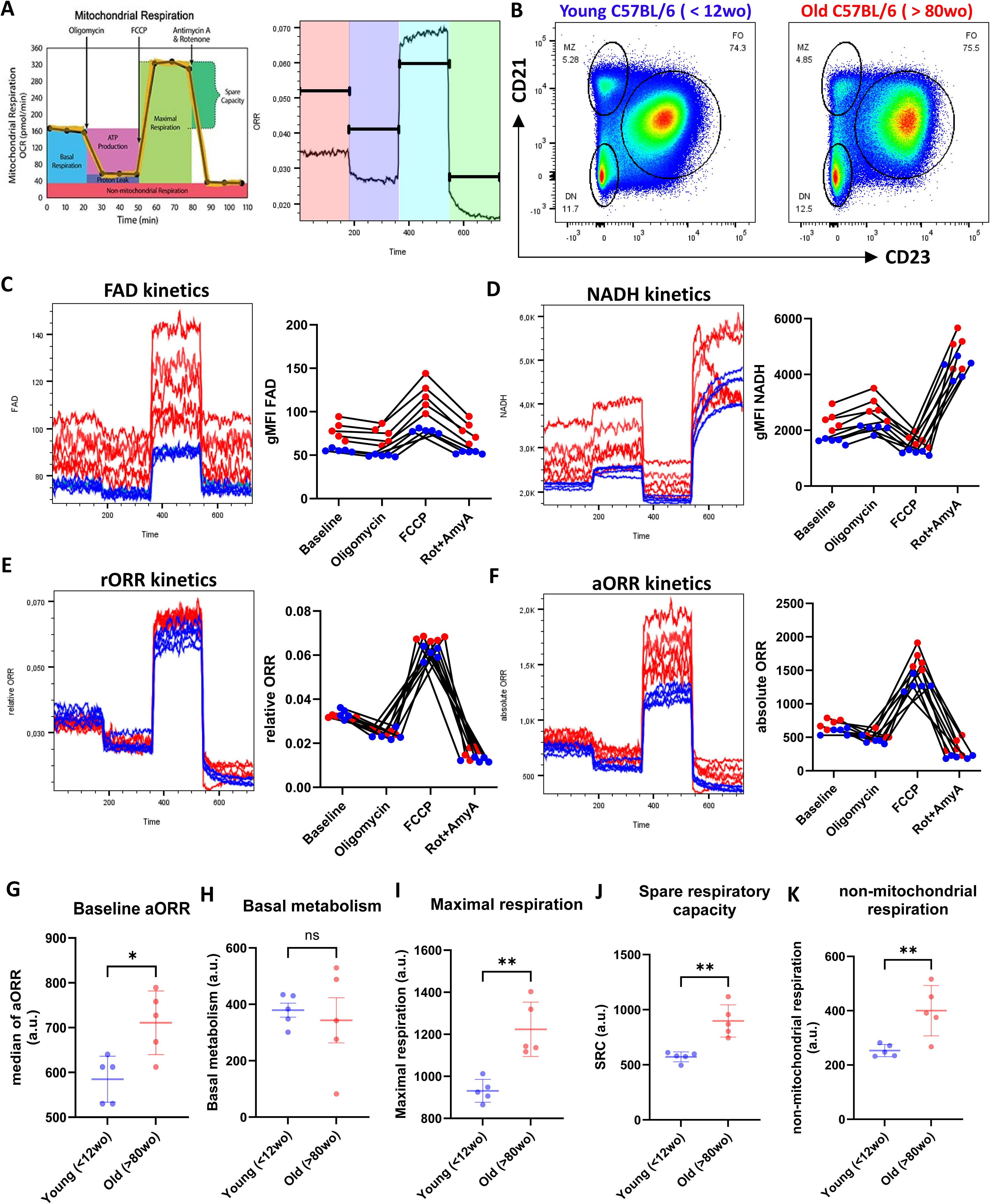
Autofluorescence-based redox profiling of ABC-like DN B cells reveals age-associated metabolic alterations between young and old wild-type C57BL/6 mice. NADH and flavin adenine dinucleotide (FAD) autofluorescence were measured to assess redox balance and mitochondrial function in CD21⁻CD23⁻ double-negative (DN) B cells isolated from young and aged wild-type C57BL/6 mice. ABC-like cells were identified as CD19⁺B220⁺CD21⁻CD23⁻ DN B cells. **(A)** Seahorse mitochondrial stress test workflow applied to DN B cells. **B)** Representative plot of splenic B2 B cell subsets, gated from CD19^+^B220^+^ B cells, from young and aged C57BL/6 wild-type mice **(C)** Real-time FAD fluorescence kinetics during mitochondrial stress testing. **(D)** Real-time NADH fluorescence kinetics during mitochondrial stress testing. **(E)** Relative optical redox ratio (rORR) kinetics during mitochondrial perturbation. **(F)** SSCA-adjusted optical redox ratio kinetics (aORR). **(G)** Baseline autofluorescence-derived adjusted optical redox ratio (aORR) at steady-state. **(H)** Basal metabolism calculated as the change in aORR (ΔaORR) between baseline conditions and following rotenone/antimycin A treatment. **(I)** Maximal respiration, calculated as median MFI of aORR following FCCP addition. **(J)** Spare respiratory capacity calculated as the ΔaORR between maximum aORR after FCCP and following rotenone/antimycin A treatment. **(K)** Non-mitochondrial respiration is calculated by the medial aORR after addition of rotenone A and antimycin. Data in graphs are shown as the mean ± SEM and analyzed using two-tailed unpaired Student’s t test. *P < 0.05, **P < 0.01, ***P < 0.001, ****P< 0.0001. Each dot represents one mouse (n = 5).

Assessment of mitochondrial redox parameters revealed that aged wild-type (>80-week-old) male C57BL/6 mice develop a hypermetabolic B-cell phenotype, predominantly driven by CD21⁻CD23⁻CD19⁺B220⁺ DN B cells **(Fig. 15A-B & Supplementary Figure S6 B-E)**. This phenotype was characterized by increased levels of both of the electron carriers NADH and FAD **(Fig. 15C-D)**, resulting in an elevated absolute optical redox rate 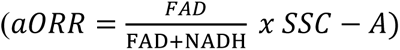, while, however, the normalized relative optical redox ratio 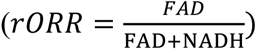 remained unchanged **(Fig. 15E-H)**. These findings indicate that the increased metabolic output reflects an expansion of overall mitochondrial mass (mitogenesis). Consistent with this interpretation, aged DN B cells displayed increased maximal respiration, enhanced spare respiratory capacity, as well as elevated non-mitochondrial FAD oxidation following rotenone/Antimycin A treatment, suggesting mobilization of alternative non-mitochondrial modes for energy productions, for instance peroxisomal fatty acid oxidation, in line with prior SEAHORSE-based studies **(Frasca et al., 2021)**. Altogether, these suggest increased metabolite consumption and energy-intensive processes, accompanied by a potential increase in ROS-generating electron transport chain leakage **(Fig. 15G-K),** as suggested by previous reports **(Ramirez De Oleo et al., 2023)**. However, our redox analysis did not show any signs of proton leak. Interestingly, analysis of the relative maximal respiratory capacity (Δ_rORR f_ollowing FCCP stimulation versus baseline) revealed differences between young and aged B cells, suggesting, besides mitogenesis, potential alterations in mitochondrial organization, architecture, or dynamics, including processes such as mitochondrial fusion or remodelling **(Fig. 15E)**. Thus, although aging-associated B cells exhibit increased metabolic capacity, this increase may reflect changes in mitochondrial quantity and network structure rather than intrinsic enhancement of mitochondrial efficiency.

## 4. Discussion

Age-associated B cells (ABCs) comprise an activated non-typical memory B cell population that accumulates during chronological aging, chronic infections, and autoimmune disorders **(Cancro, 2020; Hassani et al., 2026; Mouat et al., 2021, 2022; Mouat & Horwitz, 2022)**. Till date, ABCs have been described in numerous autoimmune diseases besides SLE, such as rheumatoid arthritis, multiple sclerosis, Sjögren’s syndrome, common variable immunodeficiency (CVID) with autoimmune complications, type I pancreatitis, Crohn’s disease and others **(Mouat et al., 2022)**. ABCs also expand in infections such as malaria, gammaherpesvirus infection, Epstein-Barr virus or SARS-CoV-2 infection, through the promotion of a type 1, interferon-rich cytokine milieu **(Mouat et al., 2021; Mouat & Horwitz, 2022)**. In the context of aging, ABC accumulation mirrors a progressive increase in systemic and localized pro-inflammatory cytokines that shape a situation described as “inflammaging” **(Cancro, 2020; Frasca et al., 2020)**. Functionally, ABCs are uniquely refractory to conventional B cell receptor (BCR) ligation alone but proliferate more robustly to endosomal nucleic acid–sensing Toll-like receptors TLR7/8 or TLR9 signalling together with type 1 cytokines **(Naradikian et al., 2016; Song et al., 2023).** However, combinatory BCR and TLR stimulation yields synergistic mutually reinforcing effects by promoting a burst of divisions **(Cancro, 2020; Naradikian et al., 2016)**.

Single-cell RNA sequencing (scRNA-seq) offers an exceptionally powerful tool for dissecting the cellular and molecular footprints, and functional programs of rare or niche immune populations. By integrating scRNA-seq datasets from patients with systemic lupus erythematosus (SLE), we were able to appreciate the phenotypic diversity and functional complexity of age-associated B cells (ABCs). Our analyses demonstrate that ABCs constitute a heterogeneous compartment encompassing several distinct subpopulations, including CD68^+^ ABCs and NKG7^+^ ABCs, characterized by transcriptional signatures associated with phagocytic/lysosomal activity and cytotoxic functions, respectively. In parallel, we observed expansion of the AREG^+^ ABC cluster. Previously, one study showed that amphiregulin, an EGF-like ligand, can be produced and secreted by B cells in the context of rheumatoid arthritis **(Mahendra et al., 2019)**. Upon binding to EGFR, it initiates processes related to immune response resolution, tissue homeostasis maintenance and tissue damage repair, and is most commonly associated with regulatory T cell **(Ehnold et al., 2025; N. Li et al., 2026)**.

Notably, STAT3 emerged as a potential upstream regulator of NKG7 expression, aligning with its established role in the development of cytotoxic natural killer T-cell populations **(Cacalano, 2016; Masle-Farquhar et al., 2022; Q. Zheng et al., 2024)**. Different cytokine pathways converge on STAT3, in particular IL6, which has been found to be enriched among the ABC subclusters **(Dai et al., 2024)**. IL6 has been implicated in SLE pathogenesis and other SLE-like diseases (e.g. Idiopathic multicentric Castleman disease), with regard to enabling the production of lupogenic anti-nuclear antibodies (ANAs) **(Uhlfelder et al., 2026)**, while IL6 knockout or neutralization with anti-IL6/-IL6R has shown benefits in the MRL.lpr model, particularly in ameliorating glomerulonephritis **(Kiberd, 1993; Reynolds et al., 2024)**. However, with regard to the potential IL6 contribution in ABC generation, heterozygous and homozygous IL6 knock-out MRL.lpr mice, still possess similar amounts of ABCs, with no alterations in splenic B2 B cell composition **(Supplementary Figure S7)**. Thus, the role of IL6 in ABC cell formation remains elusive, however, in need of further investigation.

Comprehensive transcriptomics-based immunometabolic profiling revealed pronounced metabolic heterogeneity across B-cell populations, including ABCs. Mapping glycolytic and oxidative phosphorylation (OXPHOS) activity uncovered a discrete bimodal metabolic landscape consisting of normometabolic and hypermetabolic cellular states, suggesting that metabolic changes are not uniform across all B cells. In addition, both conventional B cells and ABCs exhibited transcriptional signatures associated with cholesterol and fatty acid metabolism. ABCs demonstrated substantially greater metabolic activity than the rest of B cells. Virtually all major bioenergetic pathways were upregulated, including ketone body metabolism, pyruvate and tricarboxylic acid cycle pathways, the NADPH-generating pentose phosphate pathway, glycolysis, phospholipid metabolism, and glutathione/redox homeostasis. This metabolic configuration is more consistent with an activated effector or memory-like phenotype rather than cellular exhaustion, reflecting sustained functional engagement and continued immune responsiveness. Such metabolic adaptations could support, besides energy generation, membrane biosynthesis, biomass increase, protein production, proliferation, and protection against oxidative stress.

Progression from B cells toward the ABC state was accompanied by increased OXPHOS but also non-mitochondrial respiratory activity, potentially fuelled by lipid oxidation, in line with studies demonstrating that metabolic reprogramming facilitates B cell fate decisions towards plasma cells or memory cells **(Bradford et al., 2024; Jang et al., 2015; Lam et al., 2016; Price et al., 2018)**. A recent study elucidated a circuit between acetylcholine-secreting follicular reticular cells and autoreactive lupogenic B cells, wherein the former instigates CD36-mediated lipid uptake which in turn potentiated oxidative phosphorylation in autoreactive B cells **(Zeng et al., 2023)**. From the hdWGCNA analysis we detected signatures of acetylation-driven epigenetic reprogramming. Inspired by previous studies on RORγt^+^ Treg development by the Berod group **(Stüve et al., 2025),** we could postulate a metabo-epigenetic corridor that mobilizes OXPHOS-generated acetyl-CoA and channels it to the nucleus to modulate the chromatin landscape and histone epigenetic marks, towards ABC development. One study has already established a link between MCT1-mediated pyruvate influx into cells to secure the sufficient acetyl-CoA for acetylation of H3K27 and subsequent epigenetic activation of AID, which catalyses class-switch recombination in activated germinal center B cells **(Chi et al., 2024)**.

Pathway analyses further revealed enrichment of multiple pro-inflammatory signalling cascades, including PI3K, JAK–STAT, TNF-α, and hypoxia-associated pathways. Particularly notable was the enrichment of cytosolic DNA-sensing, Toll-like receptor and NLRP3 inflammasome pathways within the most metabolically active CD68^+^ ABC subset. Differential expression analyses identified enhanced phagocytic and lysosomal activity, alongside aberrant lupus-associated antigen processing.

Interestingly, our in-silico drug repurposing analysis with ASGARD suggested curcumin, as a potentially useful therapeutic modality, which has been shown to repress the NLRP3 pathway, via inhibition of the cytoplasmic cathepsin B **(Gu et al., 2021).** The nucleotide-binding oligomerization domain (NOD)-like receptors are a family of intracellular alarming sensors that respond robustly by the presence of viral nucleic acids, mitochondrial DNA indicative of mitochondrial disruption, toxins, ROS and other metabolic perturbations or ionic disbalance (e.g. potassium efflux, excess calcium) **(Franchi et al., 2009; Y. Liu et al., 2014; Zhong et al., 2013).** Sensing of these danger signals (alarmins) by NLRs leads to their oligomerization into large multiprotein supercomplexes that eventually trigger effector signalling cascades, such as nuclear factor-κB and MAPK kinase signalling pathways, that control the activation of pro-inflammatory caspases to curtain and combat the noxious and/or infectious stimuli **(Franchi et al., 2009; Zhong et al., 2013)**. Within the NOD-like receptor signalling pathway, ABCs displayed increased expression of multiple inflammatory mediators and innate immune signalling components, including IL18, TNF, RANTES, CXCL8, CASP1/5, IRF7, STAT1 and various interferon-regulated genes, indicating activation of inflammasome-associated and NF-κB-dependent inflammatory programs. Interestingly the ROS-induced TXNIP pathway was also augmented **(Y. Liu et al., 2014; Yang et al., 2019)**. A recent study implicated glutaminolysis in NLRP3 activation, in which they showed that pharmacological inhibition of glutaminolysis rescued lupus nephritis in MRLlpr mice, via repression of the mTOR/P70S6K/4EBP1 and NLRP3/caspase-1/IL-1β pathways **(Zhang et al., 2022).**

Our pathway visualisation portraits **(Fig. 5-6 and Supplementary Fig. S2)**, in line with the most recent literature findings **(Brown et al., 2022; Claes et al., 2016; Dai et al., 2024; Ramirez De Oleo et al., 2023; SoRelle et al., 2025)**, help us articulate an interesting chain of events. They support a model whereby excessive uptake of apoptotic debris and endogenous apoptotic material, potentially coupled with impaired lysosomal degradation, increases exposure to self-antigens. Such chronic antigenic stimulation of engulfed/phagocytosed self-material may activate intracellular RNA/DNA sensing receptors and inflammatory pathways, such as TLR7/8, cytosolic-DNA (cGAS-STING) sensing pathways, thereby promoting autoreactivity and contributing to the pathogenic functions of ABCs (such as expansion of cytotoxic NKG7^+^ ABCs). Furthermore, B cells, after being exposed to phagocytosed or endocytosed material (e.g. DNA, histones, ribonucleoprotein material), they “onload” it to the antigen-processing and presentation apparatus with which they prime T cells, or trigger self-reactive autoantibody production. TLR7/8 are endosomal RNA-sensing receptors that are triggered by viral single-stranded RNA and binds to guanosine **(Brown et al., 2022; Vinuesa et al., 2026)**. Gain-of-function mutations in the TLR7 gene were recently shown to drive lupus pathogenesis in human patients, while insertion of TLR7 mutants in mice phenocopies the human SLE pathology **(Brown et al., 2022)**. A recent study from the Vinuesa & Shen groups **(Dai et al., 2024)**, established ZEB2 as a signature transcription factor of ABCs which bestowed upon them cytotoxic and phagocytic functionalities, as the ABCs were shown, via the flow cytometric pHRodo phagocytosis assay, to phagocytose apoptotic thymocytes **(Dai et al., 2024)**.

The enhanced cholesterol metabolism observed in ABCs prompted further investigation using the MEBOCOST pipeline, which identified active biosynthesis of farnesyl pyrophosphate (FPP). Interestingly, FPP has recently been reported to accumulate in migratory dendritic cells in lupus-prone mice, where it promotes both cell survival and germinal centre responses through mitochondrial fusion and enhanced OXPHOS activity **(Zhang et al., 2024).** In the same study, inhibition of the mevalonate pathway with simvastatin restored mitochondrial homeostasis and induced disease remission **(Zhang et al., 2024)**. Our findings raise the possibility that ABCs may similarly contribute to FPP production, further supporting the notion that these cells occupy a lipid-/cholesterol-fuelled state, which could also help in coordinating spatial migratory movements of T cells or myelocytes.

ScRNA-seq-based metabolic profiling of splenic B-cell subsets from the lupus-prone MRL.lpr model recapitulated many of the observations made in human SLE. ABCs emerged as the most metabolically active B-cell population, exhibiting greater glucose and fatty acid utilization than follicular or marginal zone B cells. Likewise, cell-fate trajectory analyses positioned ABCs immediately upstream of plasma cell differentiation, corroborating previous reports demonstrating clonal continuity between ABCs and plasma cells **(Nickerson et al., 2023)**. Despite this highly active metabolic state, ABCs exhibited reduced ribosomal signatures, reflected by lower rRNA transcript abundance in both ABCs and conventional B cells from SLE patients, which was confirmed by SCENITH by checking protein translation levels between diseased and non-diseased female *vs* male MRL.lpr mice respectively **(Fig 11)**. Lower puromycin incorporation at baseline is consistent with the diminished ribosomal activity detected in complementary scRNA-seq datasets **(Fig 11E, J)**. Together, these findings reveal a paradoxical dissociation between energy production and protein synthesis.

To investigate this apparent contradiction, gene ontology and pathway enrichment analyses were performed and identified strong signatures of ubiquitination, proteasomal activity, and proteotoxic stress reminiscent of the proteotoxicity commonly observed in neurodegenerative disorders, such as prion-disease or Parkinson’s disease **(Fig. 12B, C) (Andre & Tabrizi, 2012; Di Costanzo et al., 2026).** We postulate that sustained metabolic activation drives elevated protein production, potentially overwhelming ribosomes and intracellular protein quality-control systems, including the endoplasmic reticulum and Golgi apparatus. Such stress may promote protein misfolding and accumulation of aberrant protein aggregates, thereby compromising cellular homeostasis. In response, ABCs may initiate ubiquitin-mediated proteolytic pathways to eliminate damaged proteins and maintain viability. This negative feedback mechanism could account for the reduced protein synthesis detected by puromycin incorporation, despite evidence of heightened metabolic activity. In line with the patients’ transcriptomic data, the ABCs from the MRL.lpr scRNA-seq dataset were also enriched for pathways associated with endocytosis and phagosome formation **(Fig. 12B, C).**

Finally, to determine whether these metabolic characteristics represent targetable therapeutic vulnerabilities, we performed in silico drug repurposing analyses. This in-silico approach identified inhibitors of HMG-CoA reductase inhibitors (statins) and proteasome inhibitors, as potential candidate interventions capable of targeting both the metabolic and proteostatic dependencies of ABCs respectively. These findings suggest that attacking the energetic and protein-processing infrastructure of ABCs may represent a promising therapeutic strategy for limiting their pathogenic contribution to SLE. Triggering a metabolic-blackout could perhaps result in accumulation of toxic protein aggregates which would potentially cause ABC/B cell-cell death, hopefully leading to disease-remission.

The scientific literature reports various case-studies of off-label prescription of bortezomib in incurable SLE patients, which helped combat plasmacytosis and achieved a drop in autoantibodies levels **(Walhelm et al., 2021)**. Studies in murine models also proposed a more frontline use of bortezomib in treating SLE **(Neubert et al., 2008)**. In addition, bortezomib has approval to be clinically used in multiple myeloma, where it effectively targets neoplastic plasma cells **(Field-Smith et al., 2006; Fröhlich et al., 2011)**, and could be used to treat SLE plasmacytosis.

With regard to statins repurposing of metabolic modulators, a recent meta-analysis showed profound clinical benefits in SLE patients being prescribed statins to tackle dyslipidaemia **(Qian et al., 2026)**. Statin treatment may reduce SLE disease activity and inflammatory markers (e.g., IL6, C-reactive protein, erythrocyte sedimentation rate), potentially exerting anti-inflammatory and immunomodulatory effects **(Patiño-Martinez & Kaplan, 2025; Qian et al., 2026)**. A previous study showed that fluvastatin treatment of patients for 1 month reduced the SLE Disease Activity Index and lipid levels, oxidative stress and vascular inflammation **(Ruiz-Limon et al., 2015).** Atorvastatin has been shown to repress pathologic T cell responses by intercepting with lipid raft formation, effectively disrupting T cell receptor signalling in SLE-driving T cells **(Jury et al., 2006)**

Age-associated B cells (ABCs) were originally identified as a B-cell population that accumulates in the spleens of aged mice **(Hao et al., 2011)**. ABCs were initially described in the context of biological aging, as reflected by their nomenclature. Given this established association with aging, we incorporated an age-related dimension into our analyses to determine whether ABCs arising during physiological aging exhibit metabolic features similar to those observed in autoimmune settings. Previous work by Frasca and colleagues, proposed a triangular “meta-flammatory” relationship linking inflammaging, autoantibody production, and metabolic activity **(Frasca et al., 2020, 2021, 2022).** Previous work from the group of Daniela Frasca demonstrated that ABCs from aged male mice are metabolically more active than follicular B cells and that this hypermetabolic phenotype becomes increasingly more pronounced with age **(Frasca et al., 2021)**.

SEAHORSE requires pure pre-sorted populations, which we were unable to provide, because CD21^-^ CD23^-^ DN B cells had very low viability after FACS-sorting, which would have severely confounded our measurements. To circumvent this bottleneck, we conducted a flow cytometry-based version of SEAHORSE assay **(Bartolomé & Abramov, 2015; Abir et al., 2024; Stylianakis & Hövelmeyer, 2026)**, in which we measured the relative optical redox ratio [ORR = FAD/(NADH+FAD)], a surrogate indicator of cellular respiratory activity that reflects the balance between glycolytic and oxidative metabolic processes **(Georgakoudi et al., 2025)**, throughout the application of the SEAHORSE-mitostress test. Comparison of young and aged male mice revealed that double-negative (DN) B cells exhibited higher levels of both NADH and FAD than follicular (FO) B cells, with these differences being further amplified in aged animals (**Fig. 15 and Supplementary Fig. S6)**. Although no significant differences were detected in baseline ORR measurements, aged DN B cells displayed increased maximal respiratory capacity, indicative of enhanced mitochondrial bioenergetic potential, as well as heightened non-mitochondrial respiration, derived perhaps from peroxisomal oxidation. However, after factoring the SSC-A in the optical redox ratio, we saw an increase in the adjusted aORR values in aged DN B cells. Interpreted together, this could be reflective of increased mitochondrial number and/or remodelling alterations in mitochondrial architecture, perhaps enhanced mitochondrial fusion, which is worth investigating with cryo-electron microscopy. Consistent with this interpretation, independent assessments of mitochondrial mass and membrane potential, using MitoTracker and TMRE dyes, corroborated the presence of augmented mitochondrial fitness and activity in aged DN B cells **(Fig. 14)**.

Collectively, these findings indicate that aging is associated with progressive metabolic re-wiring of the ABC/DN B-cell compartment, characterized by enhanced mitochondrial capacity and bioenergetic activity. The convergence of these metabolic adaptations with those observed in lupus-associated ABCs raises the possibility that aging and autoimmunity engage overlapping metabolic programs that promote or facilitate the expansion and functional persistence of this B-cell subset.

Targeting the metabolic infrastructure of ABCs could potentially lead to a situation of metabolic starvation or black-out of hypermetabolic and potentially pathogenic subsets, which may lead to rescission of this population in SLE.

## 5. Study limitations

While we have been able to document metabolic reprogramming in SLE, our analysis is limited by the small sample size of the scRNA-seq datasets and absence of adequate patient metadata, such as disease score or clinical history. A more comprehensive dataset, including cases of both adult and pediatric lupus at different SLEDAI disease scores, would be profoundly value-adding. Moreover, transcriptomic-based metabolic analysis, while useful in providing insights and hints, is not a perfect substitute for proteomic and metabolomic approaches, which constitute the gold standards in immunometabolic research. However, we tried to interpret the transcriptomic data in light of the flow cytometric functional assays that look directly into internal mitochondrial workings (e.g. MitoTracker/TMRE dyes, SCENITH, NADH/FAD redox profiling). Flow cytometry-based assays also provide the advantage of simultaneous examination of various subsets in parallel. Therefore, it would be worth integrating those assays with high dimensional analysis to cross-examine and compare the metabolic properties of ABCs subsets, which different based on their activation, differentiation or regulatory status, which is not possible with the standard SEAHORSE or OROBOROS platforms. Moreover, spatial analysis would also be needed to evaluate the mitochondrial architecture and structure re-engineering that may accompany SLE pathogenesis.

The MRL.lpr, while useful in translational research and able to recapitulate many aspects of the SLE disease, such as lupus nephritis and lupus autoantibodies, does not reflect the real pathogenic factors that drive SLE in humans. Loss-of-function mutations in the Fas (lpr) gene, more accurately reflect autoimmune lymphoproliferative syndrome, rather than SLE. Other mouse models like the lupus-prone B6.NZM2410 model or the *Tlr7^Y264H^*(kika) model, that carry mutations found in SLE patients, could provide more accurate results **(Brown et al., 2022; Gulinello & Putterman, 2011; Perry et al., 2011; Rubtsov et al., 2011)**. In addition, there is a gender-confounding factor in our study, since we compared mildly-diseased male with severely-disease female MRL.lpr mice, even though previous studies have also relied on such experimental design **(Nickerson et al., 2023)**.

It is still important to investigate to what extent the metabolic reprogramming observed is driven by intrinsic changes or is influenced by the surrounding cues. Here, we haven’t distinguished between endogenous and exogenous factors that may influence the metabolic reprogramming observed in B cells and ABCs. Metabolic re-wiring could be induced by intercellular interactions between B cells and other immune cells or it could be the bystander outcome of exposure to the surrounding pro-inflammatory cytokine milieu. Or more likely, it could be the outcome of intercalating and mutually re-enforcing interactions between the B cells and their niche environment. In that regard, it would be valuable to stratify a cohort of SLE patients based on the SLEDAI score, to investigate how metabolic activity alters in correlation with patients’ disease status.

We haven’t examined how various factors, such as stimulation with TLR4, TLR7, TLR9 or cytokine receptor agonists, could differentially influence metabolic activity in B cells. Our study provides a general outline on the metabolic state of DN B cells in healthy vs SLE individuals and diseased vs non-diseased MRL.lpr mice, however incorporating a temporal dimension to visualize such changes across time and disease-progression would be more value-adding. It would be interesting to cross-examine and cross-validate these findings with more relevant models, such as the recently described kika strain, since mutations in the TLR7 gene have been shown to be SLE-driving in humans.

## Supplementary Figures (Legend description)

**Supplementary Figure S1:** Standard pre-analysis quality control workflow to exclude low quality cells (with high mitochondrial counts) and doublets.

**Supplementary Figure S2: Pathway visualization across all clusters on the UMAP, with the fast gene set enrichment analysis (fgsea) package A)** Ribosome, **B)** Antigen processing and presentation. **C)** Natural killer cell mediated cytotoxicity, **D)** Systemic Lupus erythematosus, **E)** Oxidative phosphorylation, **F)** Glycolysis_Glyconeogenesis, **G)** Lysosome, **H)** Toll-like receptor signalling pathway, **I)** Fc-gamma receptor mediated phagocytosis, **J)** NOD-like receptor signalling pathway, **K)** Proteasome, **L)** Endocytosis

**Supplementary Figure S3: Pathway visualization across the three ABC subclusters clusters on the UMAP, with the fast gene set enrichment analysis (fgsea) package A)** Ribosome, **B)** Antigen processing and presentation. **C)** Natural killer cell mediated cytotoxicity, **D)** Systemic Lupus erythematosus, **E)** Oxidative phosphorylation, **F)** Glycolysis_Glyconeogenesis, **G)** Lysosome, **H)** Toll-like receptor signalling pathway, **I)** Fc-gamma receptor mediated phagocytosis, **J)** NOD-like receptor signalling pathway, **K)** Proteasome, **L)** Endocytosis

**Supplementary Figure S4:** CellChat analysis – Ligand receptor interaction partners in PBMCs from SLE patients.

**Supplementary Figure S5. CellChat analysis of intercellular communication among splenic B cell subsets in the female MRL.lpr scRNA-seq dataset (GSE202110). (A)** CellChat analysis identifying potential interaction partners with ABCs as receiver cells**. (B)** CellChat analysis identifying potential interaction partners with ABCs as sender cells. **(C)** Visualization of predicted cell–cell interactions among ex vivo immune cell populations from young and old mice.

**Supplementary Figure S6. Autofluorescence-based redox profiling reveals metabolic differences among splenic B-cell subsets in young and aged wild-type C57BL/6 mice.** NADH and flavin adenine dinucleotide (FAD) autofluorescence were measured to assess redox balance and mitochondrial metabolic activity across splenic B2 B-cell subsets. **(A)** Flow cytometric gating strategy identifying ABC-like cells according to the Hao classification, defined as CD19⁺B220⁺CD21⁻CD23⁻ double-negative (DN) B cells. **(B–E)** Seahorse’s mitochondrial stress testing workflow (baseline → oligomycin → FCCP → rotenone/antimycin **(A)** combined with autofluorescence-based measurements of NADH and FAD levels, relative optical redox ratio, and SSCA-adjusted optical redox ratio in **(B)** total B2 B cells, **(C)** follicular **(FO)** B cells, **(D)** marginal zone (MZ) B cells, and **(E)** ABC-like DN B cells.

**Supplementary Figure S7. Transcriptional pathway activity, in vitro ABC induction, and genetic modulation of B-cell phenotypes. (A–C)** Comparative pathway activity analysis of immune and metabolic transcriptional programs, including IFN responses, NOTCH and PI3K/AKT/mTOR signalling, and MYC/E2F target programs. **(D)** Induction of T-bet⁺ ABC-like B cells from MACS-purified splenic B cells from young wild-type mice under defined stimulation conditions. Cells were stimulated with BAFF, LPS, CpG, IL-21 plus IFN-γ, CpG plus IL-21 plus IFN-γ, or anti-CD40 plus IL-4 and subsequently evaluated for T-bet expression and ABC-associated phenotypic features. **(E)** decoupleR-based analysis of STAT3 expression and predicted regulatory activity. **(F)** Distribution of splenic B-cell subsets, including B2 populations and T-bet⁺CD11c⁺ DN B cells, in female MRL.lpr mice with heterozygous or homozygous IL-6 deficiency.

### Permission to reuse and Copyright

The Mitostress-test illustration depicted on figure 15A, was retrieved from this website https://www.agilent.com/en/support/cell-analysis/mitochondrial-respiration-xf-cell-mito-stress-test, after obtaining permission.

### AI usage disclosure for text editing

To improve clarity and readability, selected sections of the manuscript were refined using ChatGPT. The tool was used solely to assist with language editing after the initial draft had been prepared by the authors and did not contribute to the study design, scientific content, data analysis, interpretation of results, or conclusions. All AI-assisted revisions were critically reviewed and approved by the authors before submission.

## Conflict of Interest

The authors declare no financial, commercial or other conflict of interest.

## Author Contributions

**ES:** Conceptualization, methodology, investigation, project administration, formal analysis, data curation, literature review, data mining, bioinformatic analysis, visualization, and writing – original draft and editing. **CB:** Investigation, formal analysis, and writing – review and editing. **JWM:** Sample provision, scientific discussion and guidance, and writing – review and editing. **NH:** Funding acquisition, resources, project administration, scientific discussion, guidance and supervision, writing – review and editing.

## Acknowledgments

We would like to thank **Dr. Myriam Meineck**, from the research group of Univ.-Prof. Dr. med. Julia Weinmann-Menke, for the isolation and provision of spleen tissue samples from MRL/lpr mice, used in this study.

We would also like to thank the **FACS Core Facility CFFC PKZI FZI** of the University Medical Center of Johannes Gutenberg University Mainz, as well as its manager, **Kristian Schuetze**, for providing access to and technical support with the **BD FACSymphony™ A5 SE**, funded by the Deutsche Forschungsgemeinschaft (DFG, German Research Foundation; INST 371/45-1 FUGB), and the **Cytek® Amnis® ImageStream®X Mark II**, funded by the DFG (INST 371/43-1 FUGB). We extend our special thanks to **Dr. rer. nat. Danielle Arnold-Schild** for her technical assistance and support during the use of the ImageStream instrument.

The re-analysis of publicly available scRNA-seq datasets was performed using the high-performance computing infrastructure **MOGON 2** and advisory services provided by Johannes Gutenberg University Mainz (hpc.uni-mainz.de). MOGON 2 is part of the Alliance for High Performance Computing in Rhineland-Palatinate (AHRP; www.ahrp.info) and the Gauss Alliance e.V. The authors gratefully acknowledge the computing resources and allocation time provided on MOGON 2.

Finally, we thank **Dr. Niklas Beumer** and **Prof. Dr. Michael Delacher** for their valuable scientific input and advice regarding the bioinformatic analysis.

## Data Availability Statement

The scRNA-seq datasets used in the present study can be found here for human PBMCs GSE156751 and GSE242615. The scRNA-sew dataset corresponding to the female MRL.lpr mouse can be found here GSE202110. All primary and raw data, together with the corresponding GraphPad Prism statistical analyses, including flow cytometry FCS files, imaging flow cytometry CIF files, and Seurat objects in RDS format generated during the bioinformatic re-analysis of scRNA-seq data, have been deposited in the Zenodo repository under the ***Digital Object Identifier (DOI)*** (…). These data and associated analysis files are available upon reasonable request.

## Notes

### Competing Interest Statement

The authors have declared no competing interest.

