## Supplementary figures and images for "Single cell mapping reveals phenotypic, immunometabolic and functional heterogeneity among age-associated B cells in systemic lupus erythematosus and aging"

### Supplementary figures

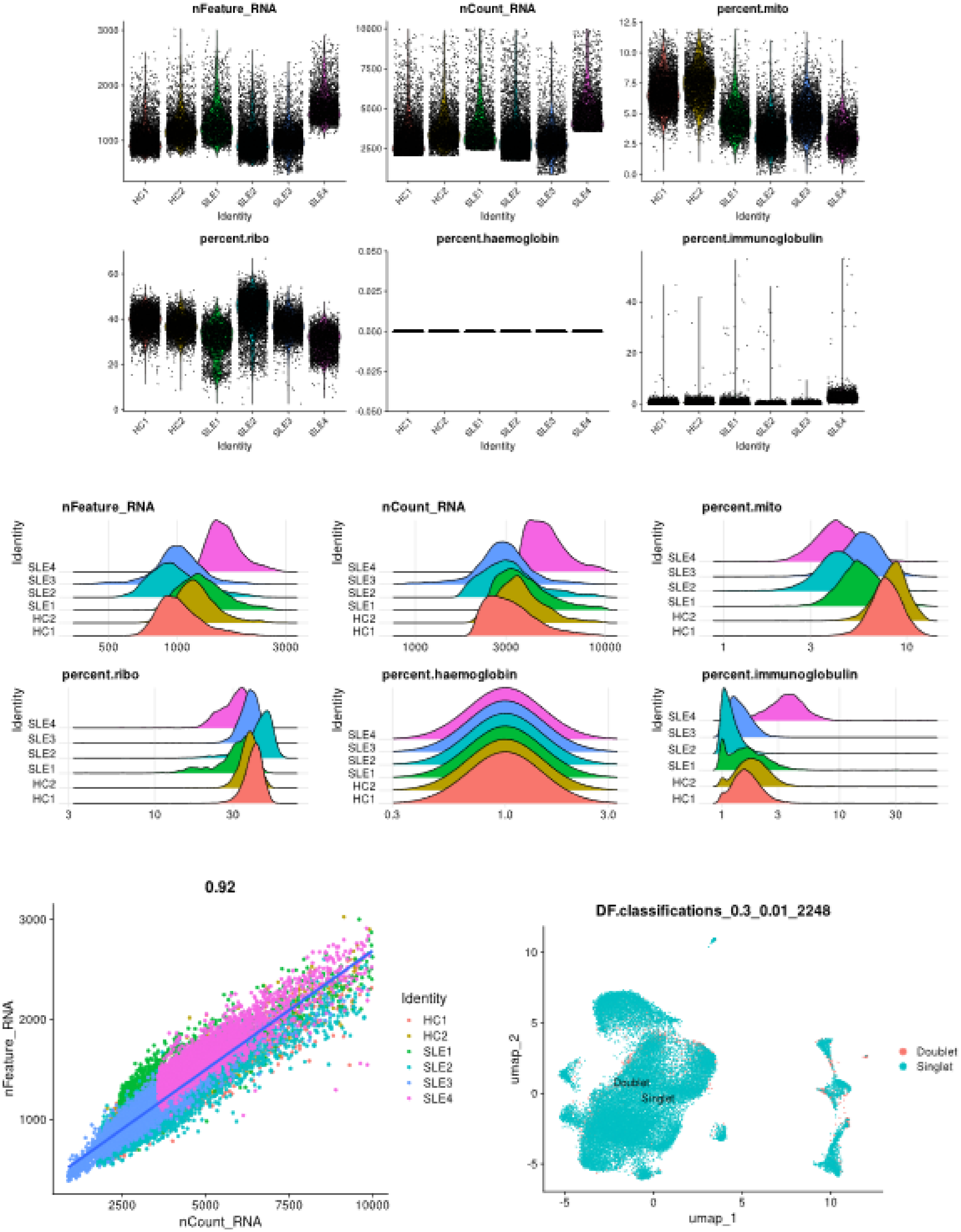

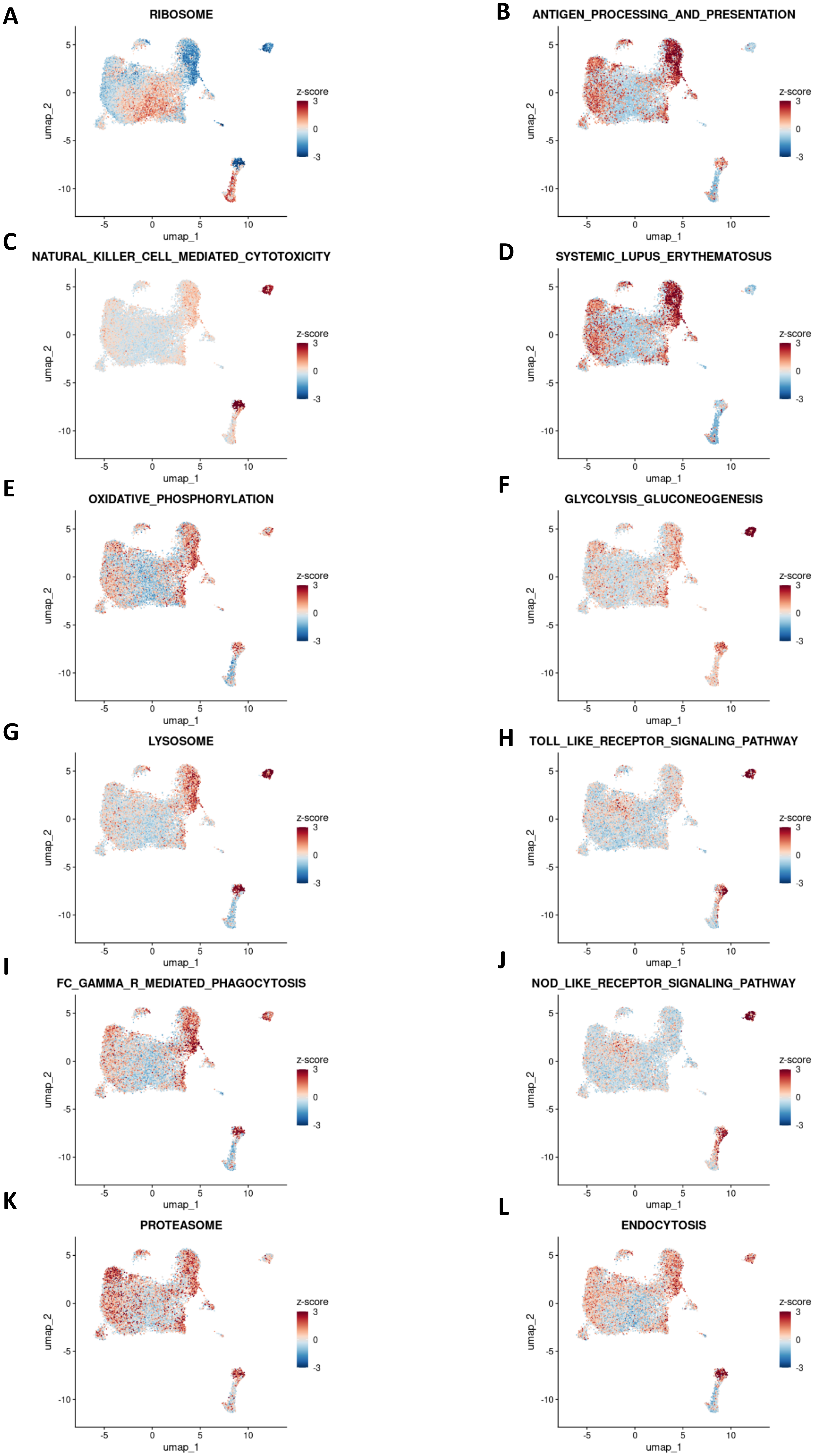

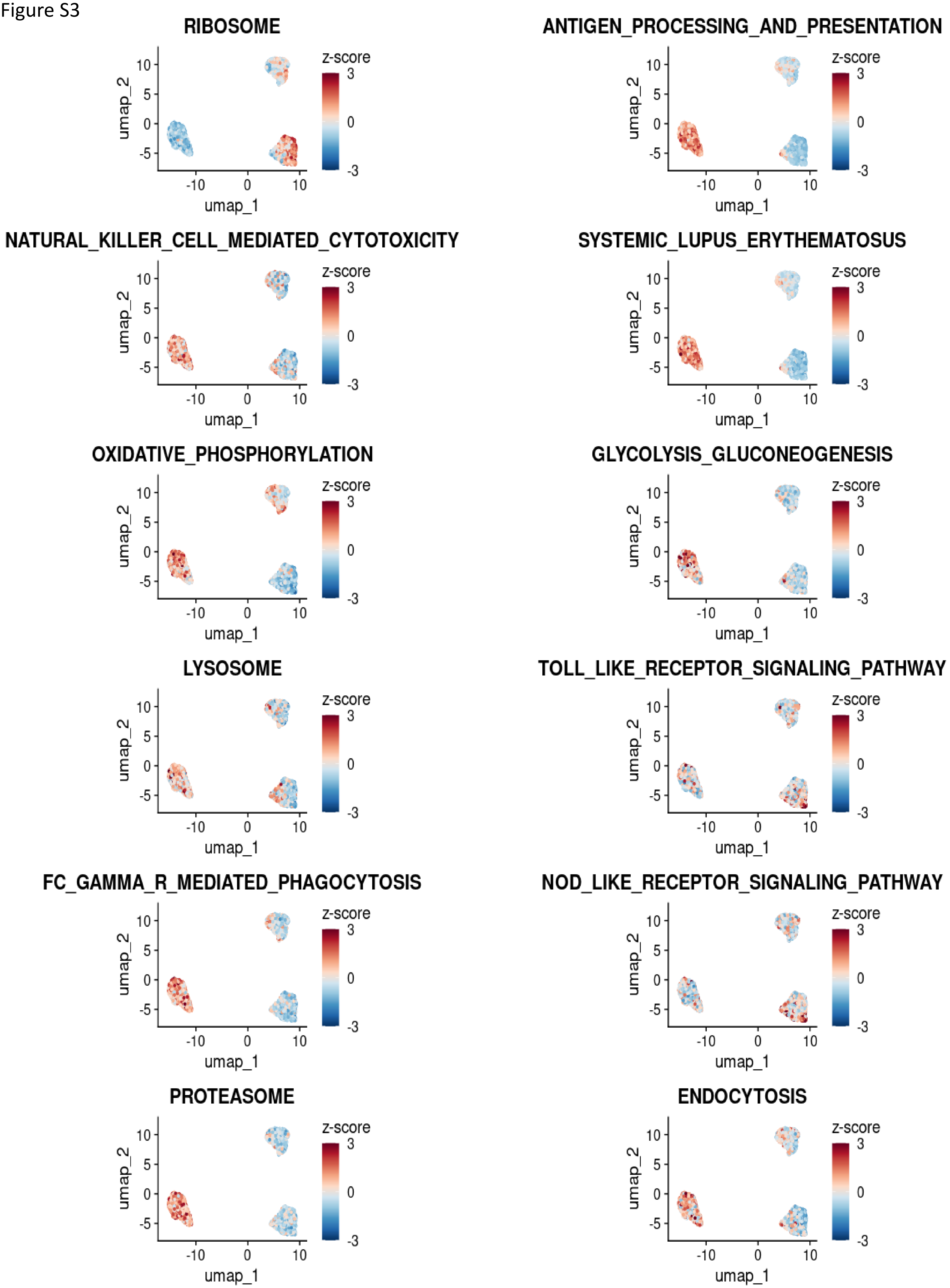

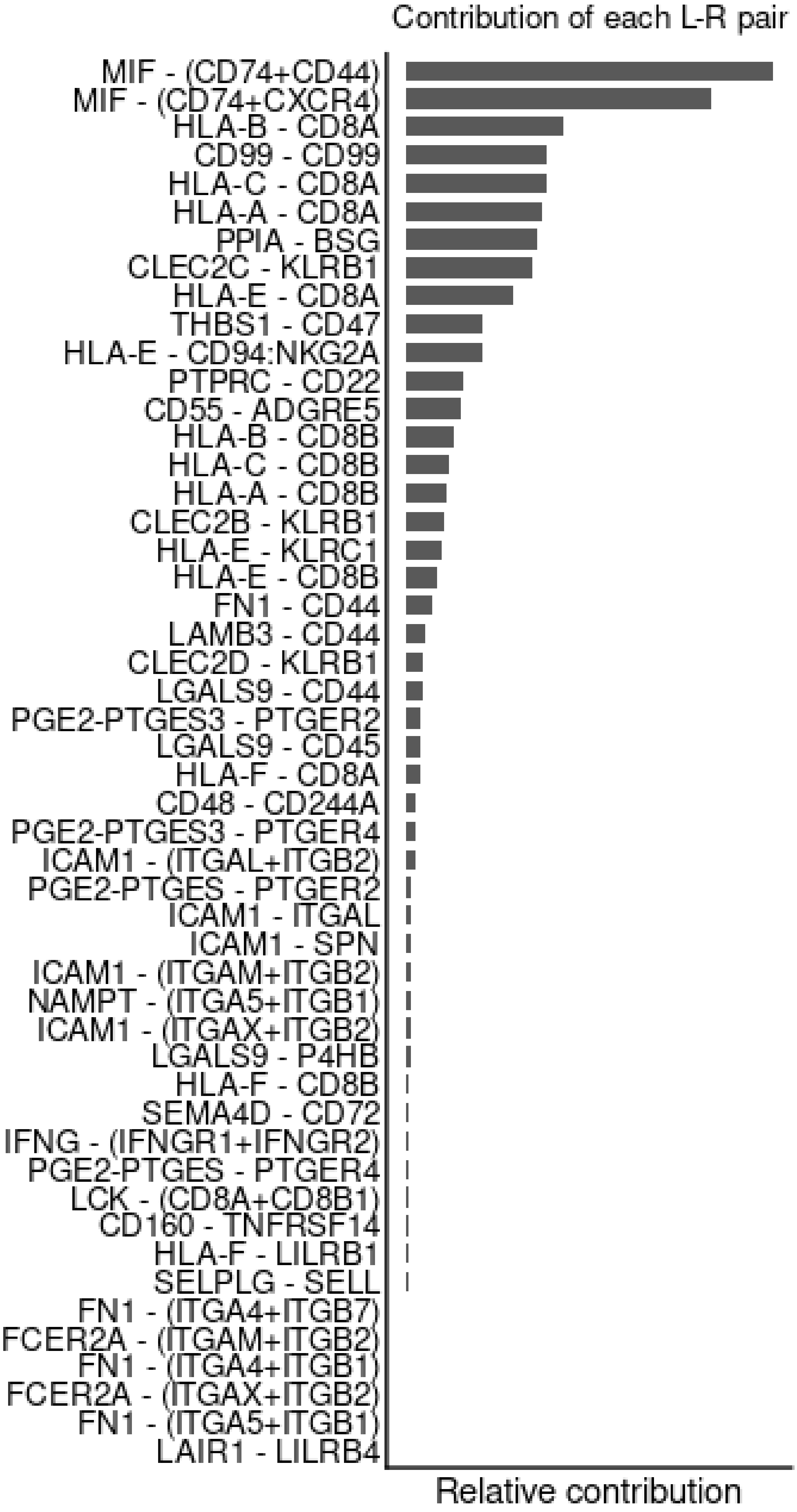

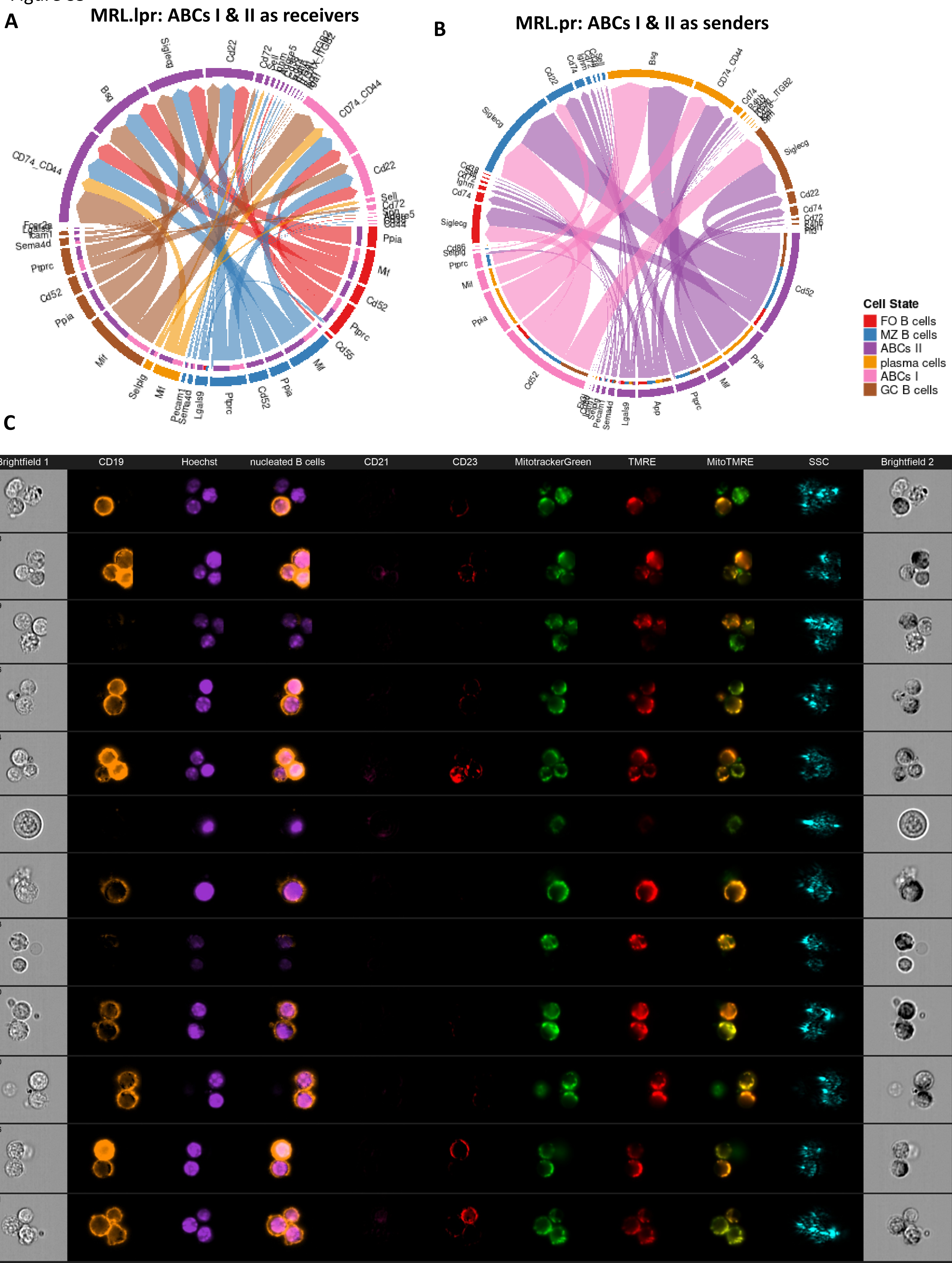

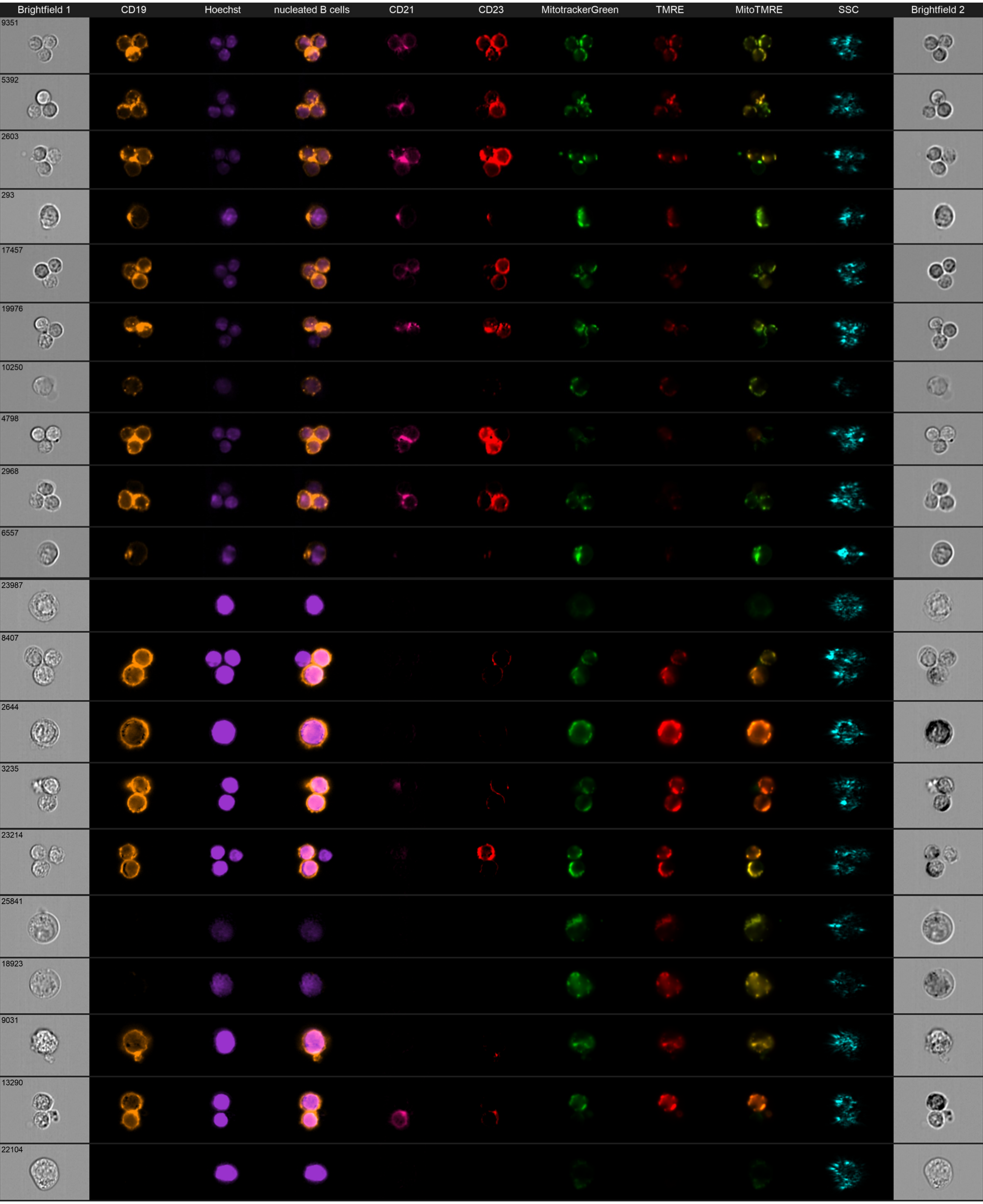

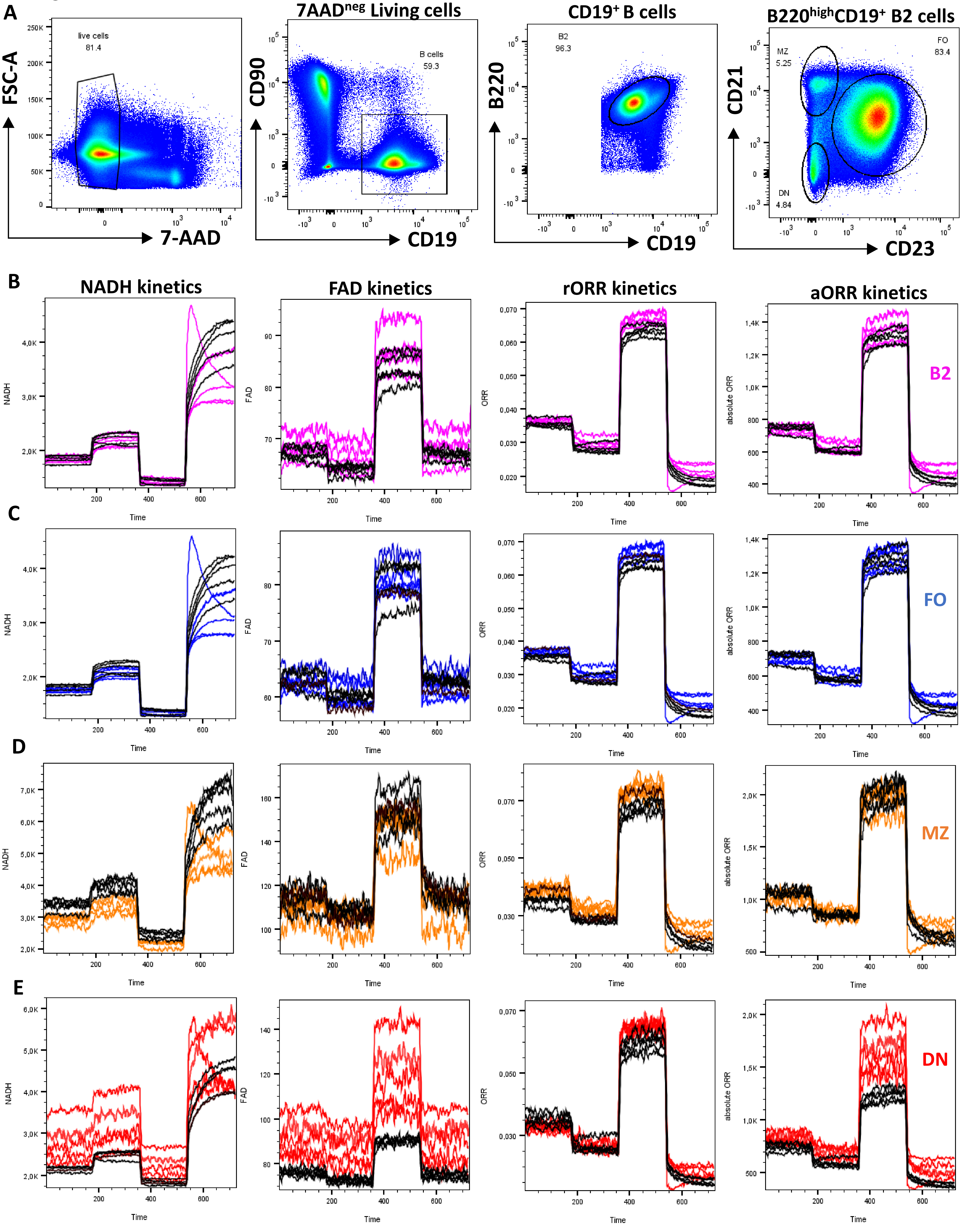

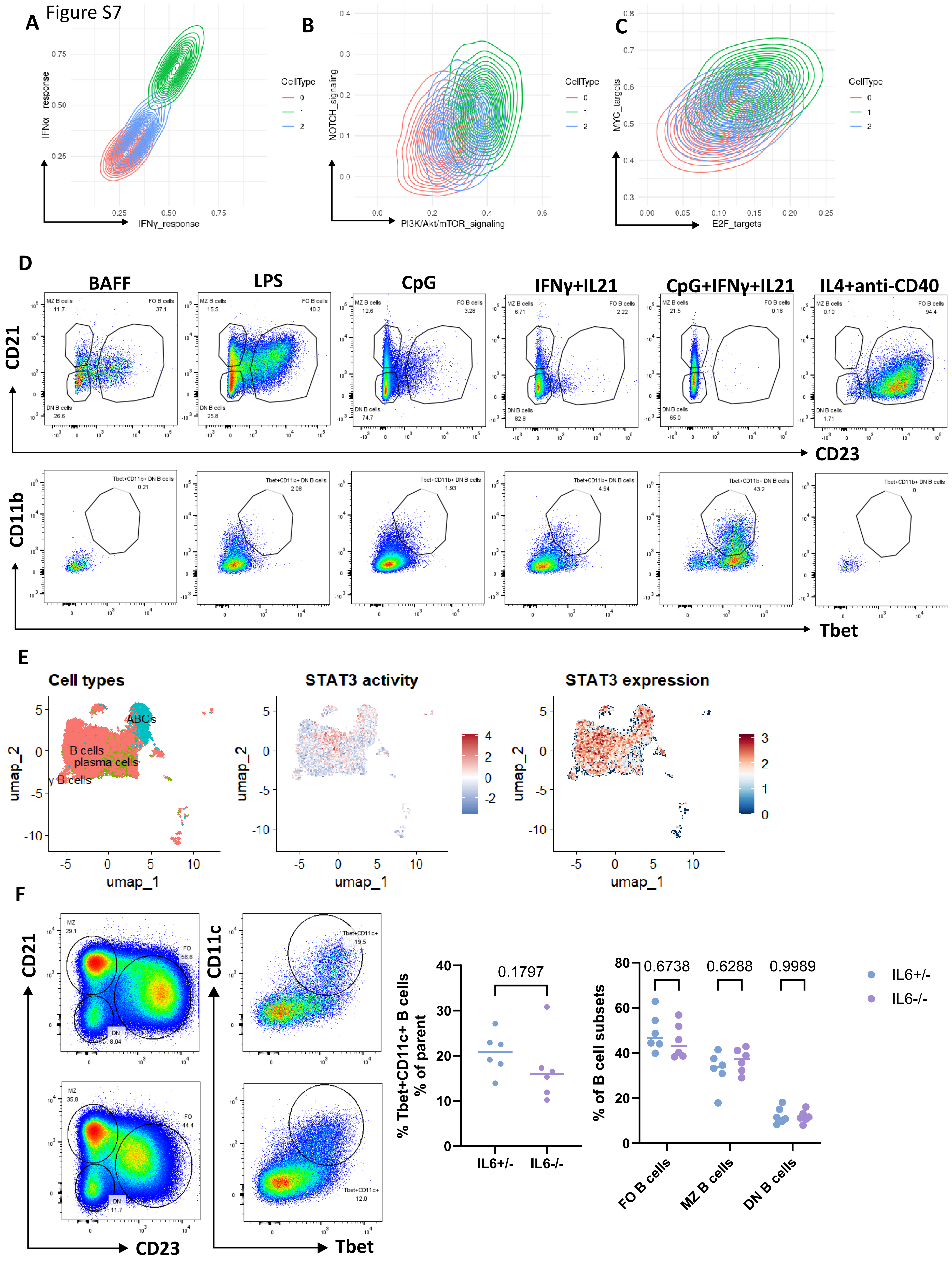
